# Druggable genome CRISPRi screen in 3D hydrogels reveals regulators of cortactin-driven actin remodeling in invading glioblastoma cells

**DOI:** 10.1101/2025.01.20.633978

**Authors:** Mufeng Hu, Anna Weldy, Isabella Lovalvo, Erin Akins, Saket Jain, Alexander Chang, Ankita Sati, Meeki Lad, Austin Lui, Akhil Rajidi, Ameya Kothekar, Erika Ding, Sanjay Kumar, Manish K. Aghi

## Abstract

To identify new therapeutic targets that limit glioblastoma (GBM) invasion, we applied druggable-genome CRISPR screens to patient-derived GBM cells in micro-dissectible biomimetic 3D hydrogel platforms that permit separation and independent analysis of core vs. invasive fractions. We identified 12 targets whose suppression limited invasion, of which ACP1 (LMW-PTP) and Aurora Kinase B (AURKB) were validated in neurosphere assays. Proximity labeling analysis identified cortactin as an ACP1- AURKB link, as cortactin undergoes serine phosphorylation by AURKB and tyrosine dephosphorylation by ACP1. Suppression of ACP1 or AURKB in culture and *in vivo* shifted the balance of cortactin phosphorylation in GBM and reduced actin polymerization and actin-cortactin co-localization. Additional biophysical analysis implicated AURKB in GBM cell adhesion and cortical stiffness, and ACP1 in resistance to mechanical stress and shape plasticity needed for 3D migration. These findings reveal a novel targetable axis that balances kinase and phosphatase activities to regulate actin polymerization during GBM invasion.

## INTRODUCTION

Glioblastoma (GBM) is the most common and lethal adult brain tumor (*1*), characterized by its robust invasive capacity (*2*). Current treatments fail to control the disease, with a median survival of just 15 months from diagnosis (*1*). Standard treatment involves maximal surgical resection followed by radiation and temozolomide chemotherapy (*3*). Unfortunately, tumor invasiveness impedes each of these three treatment modalities. First, invasiveness renders complete surgical resection impossible. Second, invasive cells migrate beyond the radiation treatment field, which is typically 2 cm beyond the enhancing edge of the tumor. Third, the escape of these invasive cells outside the area of MRI enhancement, where the blood-brain barrier (BBB) is disrupted, brings invasive GBM cells into regions where the BBB is intact, effectively protecting these invasive cells from systemic chemotherapy. These factors set the stage for the inevitable recurrence that defines GBM (*2*).

While targeting invasion has long been proposed as a strategy for treating GBM, translational progress has been slow. Two limitations of most studies to date have contributed to this lack of progress. First, existing studies have failed to recognize that infiltrating GBM cells which extend beyond the tumor edge have evolved a unique adaptive cellular machinery due to local stressors in their microenvironment. Unfortunately, these cells at the invasive tumor front are often not the ones sampled in studies analyzing banked tumor tissue, which is typically procured from the readily accessible tumor core. Second, most studies of invasion have used two-dimensional (2D) culture systems coated with a thin layer of extracellular matrix (ECM) proteins, which fail to capture the dimensionality, biomechanics, and heterogeneity of GBM invasion, as cells adherent to a 2D surface experience a different mechanical microenvironment than cells fully encapsulated in a 3D biomaterial.

To address these limitations, we have developed micro-dissectible biomimetic 3D HA-RGD hydrogels containing hyaluronic acid (HA) plus an RGD peptide sequence that enhances cell adhesion and migration by interacting with integrin receptors on cells, replicating a key feature of the GBM microenvironment (*4*). We previously leveraged the micro-dissectible nature of these devices to separate GBM cells in the implanted core from GBM cells invading the hydrogels and performed metabolomics and transcriptomics on both sets of cells to validate these devices against site-directed biopsies from patient GBMs (*4*).

In the present study, we employed a broader multi-omics approach involving a CRISPR screen targeting genes in the druggable human genome to identify targetable mediators of GBM invasion, followed by proteomics to identify key binding partners of the proteins encoded by these druggable genes (**Fig. 1**). The druggable genome refers to 2550 of the nearly 30,000 genes in the human genome that code for a protein with some precedent for binding a drug-like molecule. Most of the druggable human genome falls into seven gene families: G-protein-coupled receptors (GPCRs; n=802), serine/threonine and tyrosine protein kinases (the 519 genes comprising the human kinome), serine proteases (n=503), protein phosphatases (n=156), zinc metallopeptidases (n=106), nuclear hormone receptors (n=75), and phosphodiesterases (n=24). While less than half of these “druggable genes” have proven to be potential “drug targets” based on proven roles in human disease, of which only 120 have been targeted by marketed drugs (*5*), druggable genome screens have the added value of identifying novel mechanisms of biological processes.

**Figure 1.**
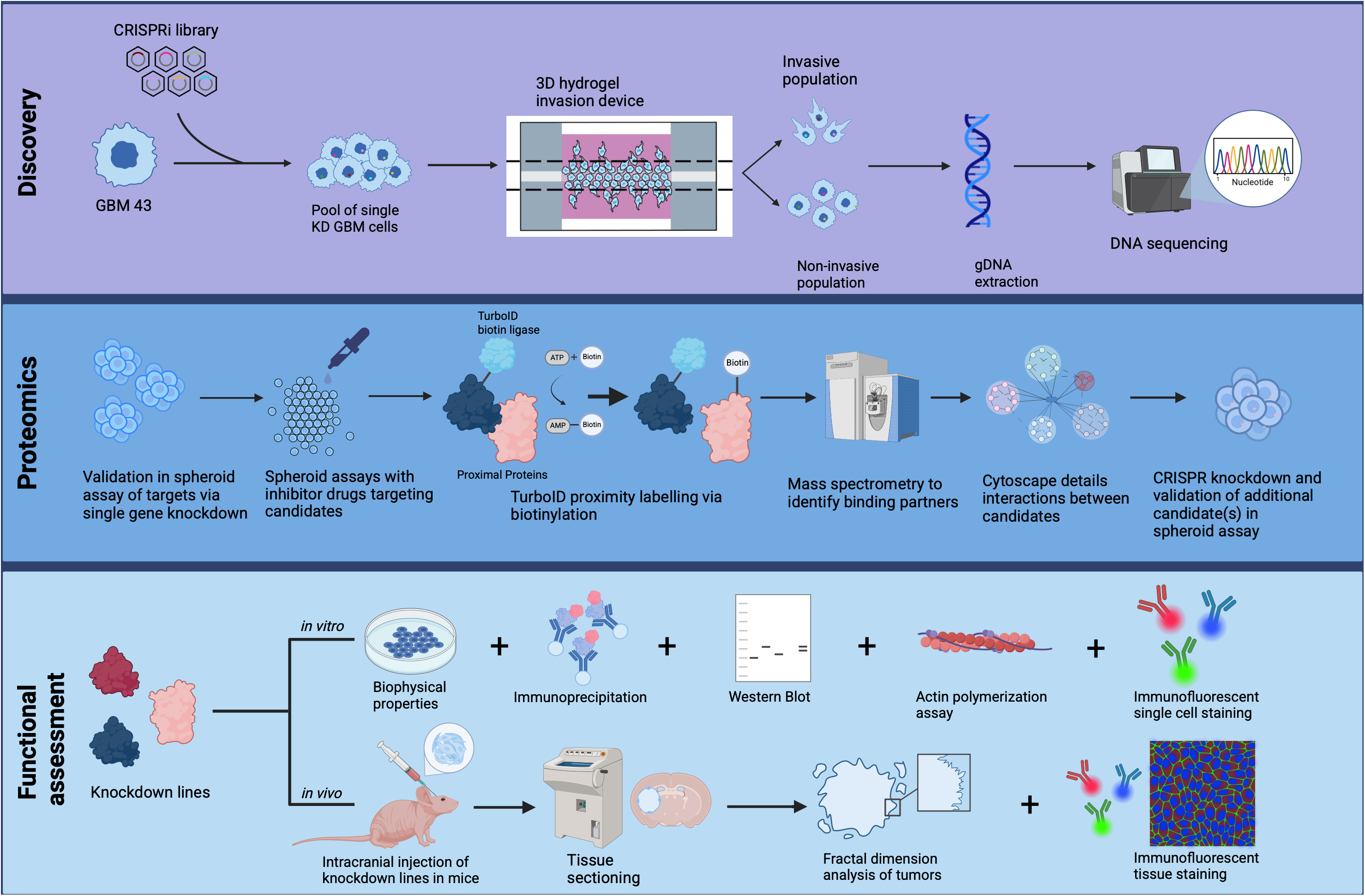
Overview of the multi-omic workflow used to identify targetable mediators of invasion in GBM. Shown are the steps involved in the discovery, proteomic, and functional assessment phases of this study. During the **discovery phase**, GBM43 cells engineered to express a CRISPRi library targeting the druggable human genome are allowed to invade 3D HA-RGD devices, after which the devices are disassembled and invasive vs. core cells are sequenced to identify sgRNAs enriched in the core cells. During the **proteomic phase**, after validating candidate sgRNAs emerging from the druggable genome screen via spheroid assays, proximity labeling assays are used to identify binding partners of the proteins encoded by the genes targeted by these sgRNAs. The binding partners can then be targeted in follow-up CRISPRi studies. During **the functional assessment phase**, functional studies were performed *in vitro*, in culture, and *in vivo* on candidates emerging from the discovery and proteomic phases.

In performing this screen, we identified the tyrosine phosphatase Acid Phosphatase 1 (ACP1; LMW-PTP) and the serine/threonine kinase Aurora Kinase B (AURKB) as druggable mediators of GBM invasion. Proximity labeling and proteomics then revealed these two proteins to share a common target: cortactin, an actin-binding protein whose phosphorylation state regulates the formation of lamellipodia. This approach bridges the gap between experimental models and clinical realities, advancing our understanding of GBM invasion and paving the way for the development of more effective therapies targeting this process.

## RESULTS

### CRISPRi screen of the druggable genome identifies novel mediators of GBM invasion

To identify druggable mediators of GBM cell invasion in an unbiased manner, we performed a CRISPRi knockdown screen with a 13,250 sgRNA library targeting the 2550 genes in the druggable human (**Supplemental Table 1**). GBM43 cells expressing dCas9-KRAB-MeCP2 and the sgRNA library were seeded in 3D HA-RGD hydrogels (n=6) and cultured for 28 days. Afterwards, devices were disassembled and micro- dissected to isolate invasive and core cells for DNA sequencing (**Supplemental Figs. 1a-b**). sgRNAs enriched in the core relative to the invasive fraction (indicating genes whose knockdown disrupted invasion) and in the invasive fraction compared to the core (indicating genes whose knockdown enabled invasion) were scored based on their abundance compared to non-targeting sgRNAs in the library **(Supplemental Table 2).** Of the 2550 genes screened, there were 12 genes with significant enrichment of targeting sgRNAs in the core (**Fig. 2a; Supplemental Table 2**): (1) RIO Kinase 2 (*RIOK2*; avg. phenotype of strongest 3=-4.6; P=0.002), an ATPase that contributes to cell growth and protein synthesis in oral squamous cell carcinoma (*6*); (2) 6- phosphofructokinase (*PFKL*; avg. phenotype of strongest 3=-3.56; P=0.004), a glycolytic enzyme that regulates lipolysis (*7*); (3) superoxide dismutase type 1 (*SOD1*; avg. phenotype of strongest 3=-2.11; P=0.0003), an antioxidant enzyme that promotes proliferation and invasion in lung cancer (*8*); (4–5) NADH:Ubiquinone Oxidoreductase Subunit A2 (*NDUFA2*; avg. phenotype of strongest 3=-2.1; P=0.002) and NADH:Ubiquinone Oxidoreductase Core Subunit V2 (*NDUFV2*; avg. phenotype of strongest 3=-1.88; P=0.0006), which are part of the electron transport chain; and (6) Phenylalanyl-tRNA synthetase beta chain (*FARSB*; avg. phenotype of strongest 3=- 3.97; P=0.003), which facilitates hepatocellular carcinoma progression by activating the mTORC1 signaling pathway (*9*); (7) TTK (avg. phenotype of strongest 3=-6.6; P=0.04), a protein kinase implicated in breast cancer metastasis (*10*); (8) Aurora Kinase B (*AURKB*; avg. phenotype of strongest 3=-3.66; P=0.002), which mediates chemoresistance in GBM (*11*); (9) acid phosphatase 1 (*ACP1*; avg. phenotype of strongest 3=-2.98; P=0.008), which mediates chemotherapy resistance and migration in colorectal cancer cells (*12*); (10) ATP6V1B2 (avg. phenotype of strongest 3=-3.6; P=0.008), which mediates intracellular organelle acidification without previously defined oncologic roles; (11) lipoic acid synthetase (*LIAS*; avg. phenotype of strongest 3=-2.6; P=0.008), a metabolic gene with a potential role in the response to apoptosis in lung adenocarcinoma cells (*13*); and (12) *GFER* (avg. phenotype of strongest 3=-3.3; P=0.001), a gene implicated in the human disulfide relay system in mitochondria without known oncologic function.

**Figure 2.**
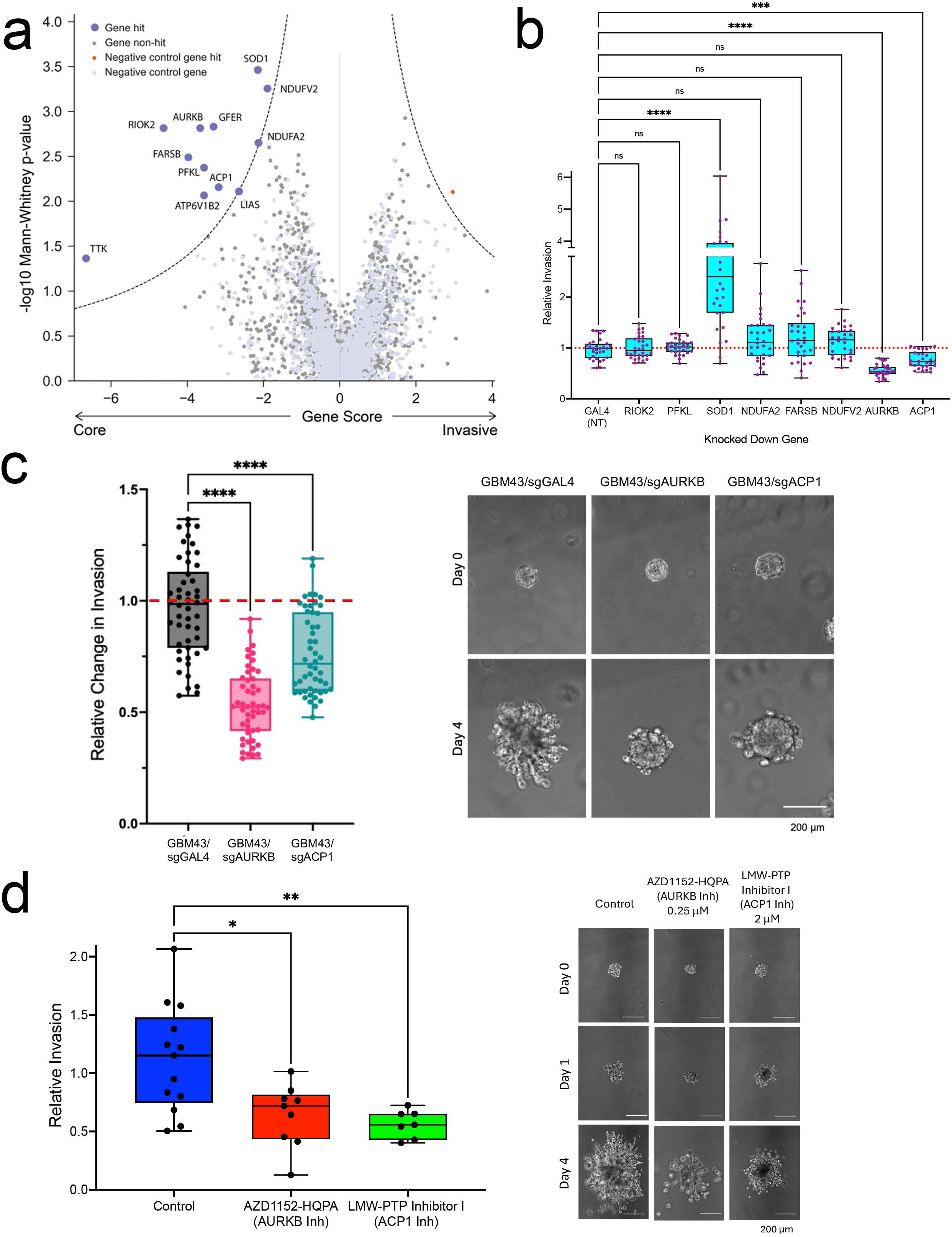
CRISPRi screen identifies druggable genes mediating GBM invasion in 3D hydrogels. **(a)** Volcano plot displaying statistical significance (y-axis) vs. magnitude of enrichment of sgRNAs for druggable genes in the invasive front vs. core (x-axis) of GBM43 cells expressing a sgRNA library targeting the druggable genome before invading 3D hydrogels. Dashed lines represent the cutoff for hit genes (false discovery rate [FDR]=0.01), with genes to the left of the dashed lines representing genes whose sgRNAs were enriched in the invasive front relative to the core. **(b)** Quantification of spheroid invasion assays of eight single gene CRISPRi knockdown GBM43 cell lines selected from CRISPR screen hits compared with control cells expressing dCas9 and sgRNA targeting GAL4 (non-targeting control=NT) (n=30 spheres/group from 3 independent experiments). **(c)** Repeat quantification with a larger sample size and representative images of spheroid invasion assays of GBM43 cells with CRISPRi knockdown of *AURKB* and *ACP1*, the only two screen hits whose CRISPRi targeting slowed GBM43 invasion. (n=48-50 spheres/group from 3 independent experiments). 10x, scale bar=200 μm. **(d)** Quantification and representative images of spheroid invasion assays of GBM43 cells treated with AURKB inhibitor AZD1152-HQPA (barasertib) or ACP1 inhibitor LMW-PTP Inhibitor I (MLS-0322825 cmpd 23), both of which slowed spheroid invasion (P=0.01 AZD1152-HQPA, P=0.006 LMW-PTP inhibitor; n=7-13 spheres/group from 3 independent experiments). 10x, scale bar=200 μm.

We then performed single-gene knockdowns on these genes to validate their contributions to slowing invasion. Substantive knockdown was achieved for 8 of the 12 candidates using single gene-targeting CRISPRi **(Supplemental Fig. 1c).** Compared to control GBM43 cells expressing dCas9, only two of eight knockdown cell lines (those targeting AURKB or ACP1) exhibited decreased invasion in spheroid assays performed in HA-RGD hydrogels (P<0.001; **Fig. 2b**).

### Validation of druggable genome CRISPRi hits with pharmacologic targeting

We then assessed the effect of pharmacologically inhibiting the proteins encoded by the two genes emerging from the druggable genome screen on GBM43 spheroid invasion: AURKB and ACP1. We chose two agents that have been investigated as anticancer agents to treat hematologic malignancies (*14, 15*): AZD1152-HQPA (barasertib), a ligand with 1000-fold selectivity for Aurora Kinase B over A (*16*), and LMW-PTP Inhibitor I (MLS-0322825 cmpd 23), an inhibitor of ACP1 with over 50-fold preference for ACP1 over a large panel of over protein tyrosine phosphatases (PTPs) (*17*). Treatment of GBM43 spheroids with AZD1152-HQPA or LMW-PTP Inhibitor I at the highest non-toxic concentrations (**Supp.** Fig. 1d) slowed invasion (P=0.01 AZD1152, P=0.006 LMW-PTP inhibitor I, **Fig. 2c**), with combined treatment of GBM43 spheroids with both agents slowing invasion even more than either agent alone (**Supp.** Fig. 1e). These findings were corroborated in two additional GBM PDX lines (GBM102 and GBM28), as treating these two lines with AZD1152-HQPA or LMW-PTP Inhibitor I also slowed 3D spheroid invasion (**Supp.** Figs. 1f-g).

### Proximity Labeling Assays Implicate Cortactin as a Shared Target of both AURKB and ACP1 in GBM

To understand the mechanisms by which AURKB and ACP1 drove GBM invasion, we first looked to identify binding partners of AURKB and ACP1 that could enable these proteins to drive their morphology-associated increases in GBM invasion. We performed proximity labeling (PL) through biotinylation coupled with mass spectrometry (MS), which has emerged as a powerful unbiased technique for capturing protein-protein interactions (PPIs) within living cells. We used the Turbo-ID system (*18*) which fuses the protein of interest to a biotin ligase that uses ATP to convert biotin into biotin–AMP, a reactive intermediate that covalently labels proximal proteins. We then performed streptavidin bead pull-down of lysates from GBM43 cells expressing Turbo- ID AURKB and ACP1 fusion proteins (**Fig. 3a**), followed by MS analysis of the pulled-down biotinylated proteins. Amongst the KEGG pathways enriched for these AURKB and ACP1 binding proteins, three were relevant to cancer invasion: proteoglycans in cancer, regulation of the actin cytoskeleton, and focal adhesion (**Fig. 3b**). Volcano plots revealed that of 2,814 different proteins identified by MS, 1,689 and 1,889 bound significantly to Turbo-ID AURKB and ACP1 fusion proteins, respectively, relative to the control lysates from cells expressing a Turbo-ID cytochrome P450 fusion protein, with 1,347 proteins shared between the two (**Supp. Table 3**, **Fig. 3c**). Amongst the 1,347 proteins interacting with AURKB and ACP1, 8 were found in the three invasion-relevant KEGG pathways mentioned above: Eukaryotic translation initiation factor 4B (*EIF4B*), integrin-linked kinase (*ILK*), Rac/Cdc42 Guanine Nucleotide Exchange Factor 6 (*ARHGEF6*), Rho Guanine Nucleotide Exchange Factor 7 (*ARHGEF7*), G-protein-coupled receptor kinase interactor-1 (*GIT1*), p21 protein-activated kinase 2 (*PAK2*), thymosin-β4 (*TMSB4X*), and cortactin (*CTTN*) (**Fig. 3c**). We then refined this list by identifying which of these were upregulated in our previous RNA-seq of GBM43 cells invading 3D hydrogels (*4*), which narrowed the list down to two genes, both of which encode actin-binding proteins: *TMSB4X* and *CTTN* (**Fig. 3d**). Because thymosin- β4 (encoded by the *TMSB4X* gene) only has serine and threonine phosphorylation sites (hence only actable on by AURKB), while cortactin (encoded by the *CTTN* gene) has serine, threonine, and tyrosine phosphorylation sites (*19*) that could be acted on by both AURKB and ACP1, we focused on cortactin, an actin-binding protein with two known roles in cell morphology and migration: (1) regulating the interactions between components of adherens-type junctions and (2) organizing the actin cytoskeleton and cell adhesion structures such as lamellipodia of epithelia and cancer cells. Further rationale for focusing on cortactin emerged from *in silico* analysis performed using the open-source platform Cytoscape, a database of PPI networks, revealing that cortactin interacts with numerous proteins known to interact with AURKB and ACP1 (**Fig. 3e**), and from 3D spheroid assays in which *CTTN* knockdown (**Supp.** Fig. 2a) led to reduced GBM43 invasion (**Supp Fig. 2b**), similar to what was seen with *AURKB* or *ACP1* knockdown.

**Figure 3.**
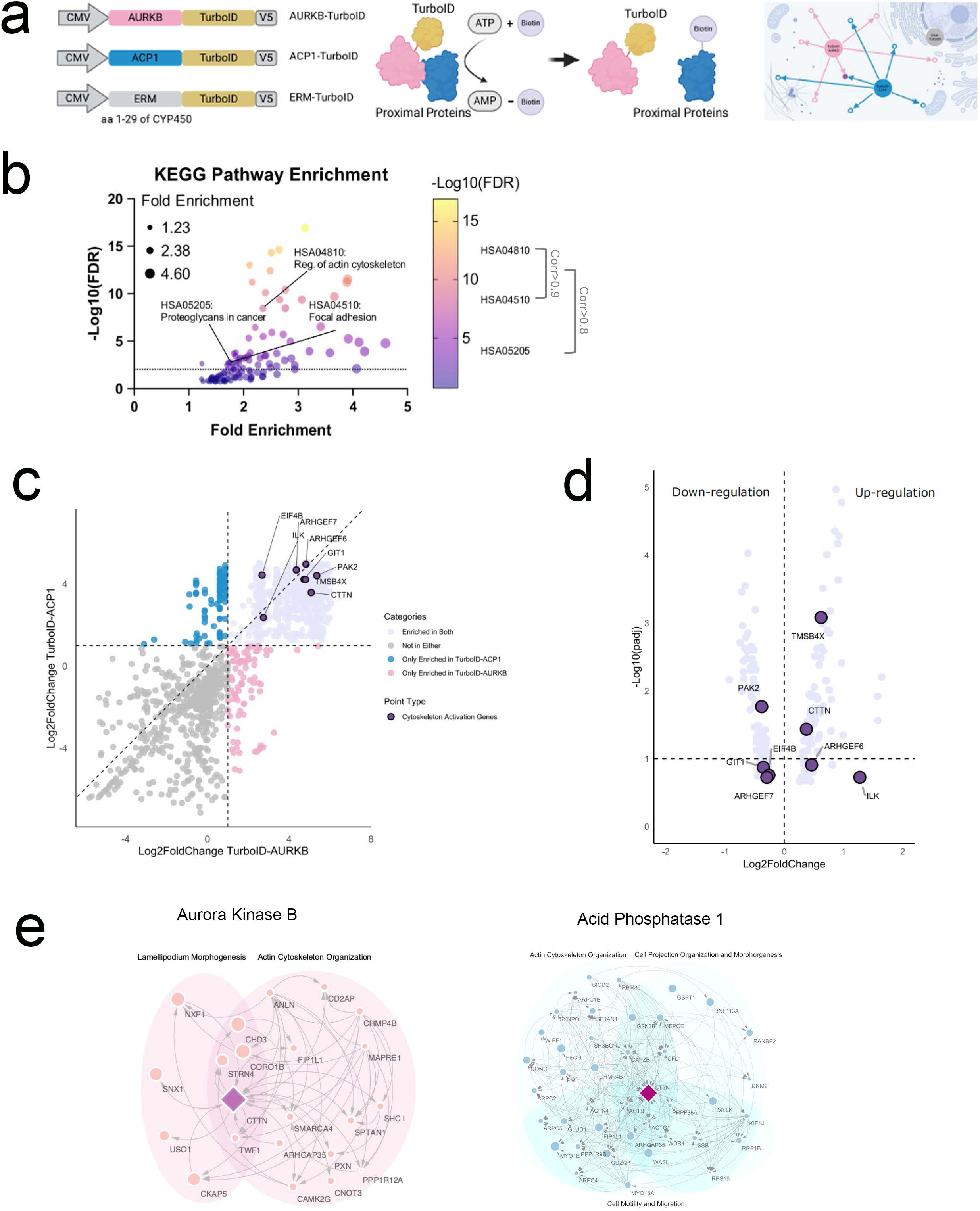
Proximity labeling identifies cortactin as a binding partner for AURKB and ACP1, and a key regulator of GBM invasion, whose expression increases in invasive GBM cells. (a) Workflow for Turbo-ID AURKB and ACP1. Plasmids were created in which AURKB, ACP1, and control sequence encoding amino acids 1-29 of cytochrome P450 were fused to the TurboID sequence, an engineered biotin ligase that uses ATP to convert biotin into biotin–AMP, a reactive intermediate that covalently labels proximal proteins. (b) Shown are KEGG pathways shared amongst proteins bound to both AURKB and ACP1 plotted based on their fold enrichment (x-axis) and -log_10_(FDR) (y-axis) with pathways related to invasion highlighted. Three of these pathways are relevant to cancer invasion and are highlighted along with the correlation between their gene sets: proteoglycans in cancer, regulation of the actin cytoskeleton, and focal adhesion. (c) Proteins bound to AURKB or ACP1 listed based on the log_2_Fold Change of their binding to AURKB relative to cytochrome P450 (x-axis) and the log_2_Fold Change of their binding to ACP1 relative to cytochrome P450 (y-axis). Proteins shaded gray did not exhibit significant binding to either protein relative to control. Proteins in cyan exhibited significant binding to ACP1 relative to control. Proteins in pink exhibited significant binding to AURKB relative to control. Proteins in light purple exhibited significant binding to AURKB and ACP1 relative to control. The eight proteins shaded dark purple and identified are part of the three invasion-related KEGG pathways highlighted in (**b**). (d) Genes for proteins that significantly bound AURKB and ACP1 are stratified according to their gene expression in the edge vs. core fractions of 3D hydrogels into which GBM43 cells invaded. (e) CellSCAPE analysis of how cortactin interacts with proteins known to interact with AURKB (*left*) and ACP1 (*right*)

### AURKB and ACP1 regulate cortactin phosphorylation in GBM

We then further interrogated the interactions between cortactin and the AURKB and ACP1 proteins. We used immunoprecipitation to confirm that cortactin bound both AURKB and ACP1 (**Fig. 4a**). GBM43 cells with *AURKB* knockdown or treated with AZD1152-HQPA exhibited decreased serine/threonine phosphorylation of cortactin (**Fig. 4b; Supp.** Fig. 3a), consistent with AURKB acting as a serine/threonine kinase on cortactin. Because ACP1 is a tyrosine phosphatase, we then investigated tyrosine phosphorylation of cortactin in GBM43 cells with *ACP1* knockdown or with LMW-PTP Inhibitor I treatment. Targeting ACP1 with knockdown or with LMW-PTP Inhibitor I led to increased tyrosine phosphorylation of cortactin (**Fig. 4c; Supp.** Fig. 3b). Together, these findings suggested that cortactin undergoes serine phosphorylation by AURKB and tyrosine dephosphorylation by ACP1. Furthermore, analysis of total cortactin levels by Western blot in these cells revealed that GBM43/sgAURKB cells had less cortactin than control GBM43/sgGAL4 cells (**Fig. 4d**), suggesting that cortactin’s phosphorylation status impacted the stability of the protein.

**Figure 4.**
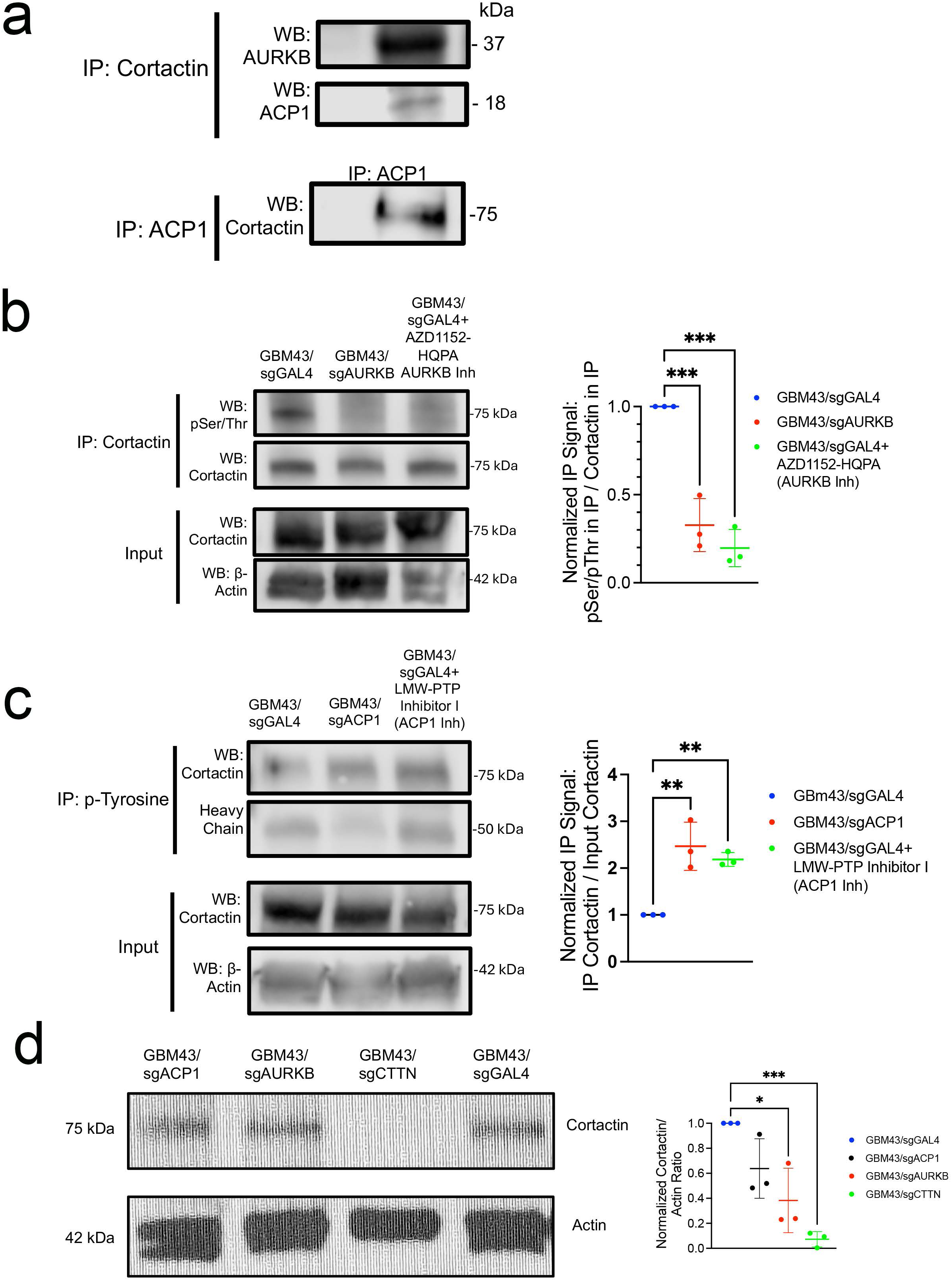
Serine phosphorylation of cortactin by AURKB and tyrosine dephosphorylation of cortactin by ACP1. **(a)** Immunoprecipitation of cortactin from GBM43 cell lysates followed by blotting for AURKB (upper row) and ACP1 (middle row), along with immunoprecipitation of ACP1 followed by blotting for cortactin (lower row) confirms cortactin binding to AURKB and ACP1. **(b)** Targeting AURKB with CRISPRi in GBM43/sgAURKB cells or via the drug AZD1152-HQPA leads to reduced cortactin serine phosphorylation, as determined by immunoprecipitation of cortactin followed by blotting for phosphorylated serine/threonine (first row) and blotting for cortactin (second row) or blotting the lysates without precipitation for cortactin (third row) or β-actin (fourth row). Quantification shown in the graph to the right was derived from band intensities in 3 technical replicates. **(c)** Targeting ACP1 with CRISPRi in GBM43/sgACP1 cells or via the drug LMW-PTP Inhibitor I leads to increased cortactin tyrosine phosphorylation, as determined by immunoprecipitation for phosphorylated tyrosine followed by blotting for cortactin (first row) and blotting for heavy chain antibody (second row), as well as blotting the non-precipitated lysates for cortactin (third row) and beta-actin (fourth row). Quantification shown in the graph to the right was derived from band intensities in 3 technical replicates. **(d)** Western blot for total cortactin levels confirms *CTTN* knockdown in GBM43/sgCTTN cells relative to GBM43/sgGAL4 control cells and also reveals decreased cortactin levels in GBM43/sgAURKB cells relative to GBM43/sgGAL4 control cells, as determined by quantification of band intensities in 3 technical replicates to generate the graph to the right. *P<0.05; **P<0.01; ***P<0.001; ****P<0.0001. Scatter dot plots show the mean (horizontal bar) with standard deviation (vertical bar).

### Cortactin Drives Actin Polymerization in GBM Lamellipodia and Associated Morphologic Changes Upon Phosphorylation

Given the role of cortactin in actin polymerization (*20*), we then investigated the impact of targeting *AURKB* or *ACP1* on actin polymerization using an *in vitro* assay performed on lysates of GBM43 cells with *AURKB* or *ACP1* knockdown. We found that AURKB (**Fig. 5a**) or ACP1 (**Fig. 5b**) knockdown or pharmacologic targeting led to reduced rates of actin polymerization (). Similarly, *CTTN* knockdown in GBM43 cells also led to reduced rates of actin polymerization (**Supp.** Fig. 4a).

**Figure 5.**
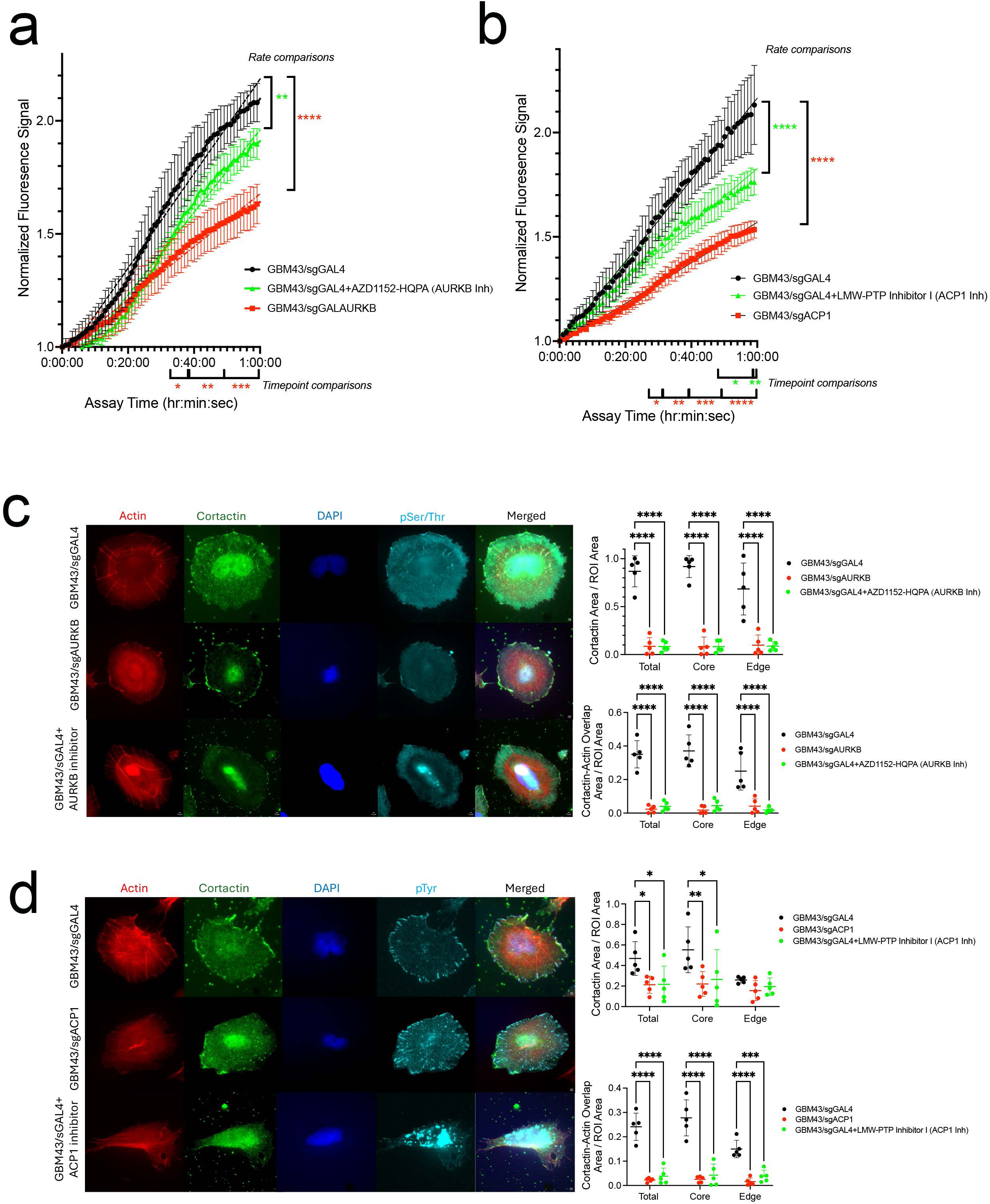
AURKB and ACP1 affect cortactin phosphorylation, altering actin polymerization, cortactin levels, and actin-cortactin overlap. **(a)** Lysates from GBM43/sgGAL4 cells (black), GBM43/sgGAL4 cells treated with AURKB inhibitor AZD1152-HQPA (green), and GBM43/sgAURKB cells (red) were used in actin polymerization assays, with lysates from GBM43/sgGAL4 cells treated with AURKB inhibitor AZD1152-HQPA (P=0.003) and GBM43/sgAURKB cells (P<0.0001) slowing the rate of actin polymerization relative to GBM43/sgGAL4 lysates (assessed by comparing the slope of the polymerization curves derived from linear regression). Differences in polymerization levels at individual timepoints occurred for GBM43/sgAURKB lysates vs. GBM43/sgGAL4 lysates as shown by red asterisks below the x-axis, differences in polymerization between lysates of GBM43/sgGAL4 cells treated with AZD1152-HQPA vs. GBM43/sgGAL4 lysates were not significant during the range of times assessed, although the differences in actin polymerization rates between the two conditions suggested they would eventually differ. n=3/group. Points represent means, error bars represent standard errors. **(b)** Lysates from GBM43/sgGAL4 cells (black), GBM43/sgGAL4 cells treated with LMW-PTP Inhibitor I to target ACP1 (green), and GBM43/sgACP1 cells (red) were used in actin polymerization assays, with lysates from GBM43/sgGAL4 cells treated with LMW-PTP Inhibitor I to target ACP1 (P<0.0001) and GBM43/sgACP1 cells (P<0.0001) slowing the rate of actin polymerization relative to GBM43/sgGAL4 lysates (assessed by comparing the slope of the polymerization curves derived from linear regression). Differences in polymerization levels at individual timepoints occurred for GBM43/sgACP1 lysates vs. GBM43/sgGAL4 lysates as shown by red asterisks below the x-axis, differences in polymerization between lysates of GBM43/sgGAL4 cells treated with LMW-PTP Inhibitor I vs. GBM43/sgGAL4 lysates at individual timepoints are shown by green asterisks below the x-axis. n=3/group. Points represent means, error bars represent standard errors. **(c)** Targeting AURKB with CRISPRi or drug (AZD1152-HQPA) in cultured GBM43 cells (n=5/group) leads to reduced cortactin levels (P<0.0001, upper graph) and reduced cortactin-actin overlap (P<0.0001; lower graph) normalized to cell area at a total intracellular level as well as when separating the cell core from cell edge. 60x magnification, scale bar: 2 μm. 60x magnification, scale bar: 2 μm. Scatter dot plots show the mean (horizontal bar) with standard deviation (vertical bar). **(d)** Targeting ACP1 with CRISPRi or drug (LMW-PTP Inhibitor I) in cultured GBM43 cells (n=5/group) leads to reduced cortactin immunostaining at a total intracellular level (P=0.042 CRISPRi and P=0.045 inhibitor) and in the cell core (P=0.006 CRISPRi and P=0.02 inhibitor), but not at the cell edge (P=0.6-0.8) where background levels of cortactin started out lower (upper graph) and reduced cortactin-actin overlap throughout the cell (P<0.0001 total, core, and edge; lower graph) normalized to cell area. 60x magnification, scale bar: 2 μm. Scatter dot plots show the mean (horizontal bar) with standard deviation (vertical bar). *P<0.05; **P<0.01; ***P<0.001; ****P<0.0001.

These effects led to morphologic changes in cultured GBM43 cells, in which *AURKB* or *ACP1* knockdown lowered the aspect ratio, a morphologic metric that is elevated in elongated spindle-like cells more adept at invasion compared to rounder cells (*21*) (P<0.0001; **Supp.** Fig. 4b). These morphologic changes led to GBM43 cells with *AURKB* or *ACP1* knockdown having smaller starting volume than control GBM43/sgGAL4 cells upon encapsulation in 3D hydrogels (**Supp.** Fig. 4c). We then investigated the roles of AURKB and ACP1 on the response of GBM cells to hyperosmotic stress caused by incubation with hypertonic polyethylene glycol (PEG) solution, which causes cells to lose intracellular water and thereby shrink. We found that GBM43 cells with *AURKB* or *ACP1* knockdown exhibited resistance to hyperosmotic stress-induced cellular compression because cells with knockdown of these two genes were already so compacted that hyperosmotic stress could not compress them any further (**Supp.** Fig. 4d), suggesting that targeting AURKB or ACP1 compromised the shape plasticity needed for rapid 3D migration (*22*).

Given the importance of cellular localization of actin and cortactin for lamellipodia formation at the edge of invading cancer cells, we then used immunofluorescent analysis of cultured cells to determine if there were regional intracellular variations in the phosphorylated cortactin changes we noted by Western blot with AURKB or ACP1 targeting in GBM43 cells (**Fig. 4c**). We found that the diminished cortactin serine phosphorylation achieved with AURKB knockdown or pharmacologic targeting were demonstratable by IHC throughout the cell core or within 3 μm of the cell edge (**Supp.** Fig. 4e). On the other hand, the increased tyrosine phosphorylation caused by ACP1 knockdown or pharmacologic targeting could be demonstrated in total throughout the cell and in the cell core, but not at the cell edge, due to a high background level of cortactin tyrosine phosphorylation at the cell edge (**Supp.** Fig. 4f). We also found that AURKB targeting by knockdown or drug inhibition reduced total cortactin levels throughout the cell in both the core and edge (**Fig. 5c**), while ACP1 targeting by knockdown or drug inhibition only reduced cortactin at the cell core, not the cell edge (**Fig. 5d**), consistent with which region experienced increased cortactin tyrosine phosphorylation with ACP1 targeting.

Given the effects of these proteins noted on actin polymerization, we then investigated the impact of AURKB and ACP1 targeting on the cellular co-localization of cortactin and actin. We found that AURKB and ACP1 targeting with knockdown or drug treatment disrupted actin-cortactin colocalization throughout the cell core and edge (**Figs. 5c-d**).

### Effects of AURKB, ACP1, and CTTN on actin remodeling and GBM invasion *in vivo*

We then investigated the impact of *AURKB*, *ACP1*, or *CTTN* knockdown on GBM invasion *in vivo*. Invasiveness *in vivo* was assessed by fractal analysis of images of tumors and their surrounding brain, yielding the fractal dimension, a numeric description of invasive tumor growth pattern as a number between 1 and 2, with higher numbers representing greater invasiveness (*4*). This method revealed that GBM43 cells with *CTTN* or *AURKB* knockdown, but not those with *ACP1* knockdown, exhibited decreased invasion *in vivo* (**Fig. 6a**). While Kaplan-Meier analysis revealed that none of the 3 knockdowns altered survival compared to GBM43/sgGAL4 control xenografts (**Fig. 6b**), with GBM43/sgACP1 and SGBM43/sgCTTN xenografts achieving smaller cross-sectional area than GBM43/sgGAL4 xenografts (**Fig. 6c**), again suggesting a benefit in targeting these genes *in vivo*, albeit a benefit that fell short of impacting survival.

**Figure 6.**
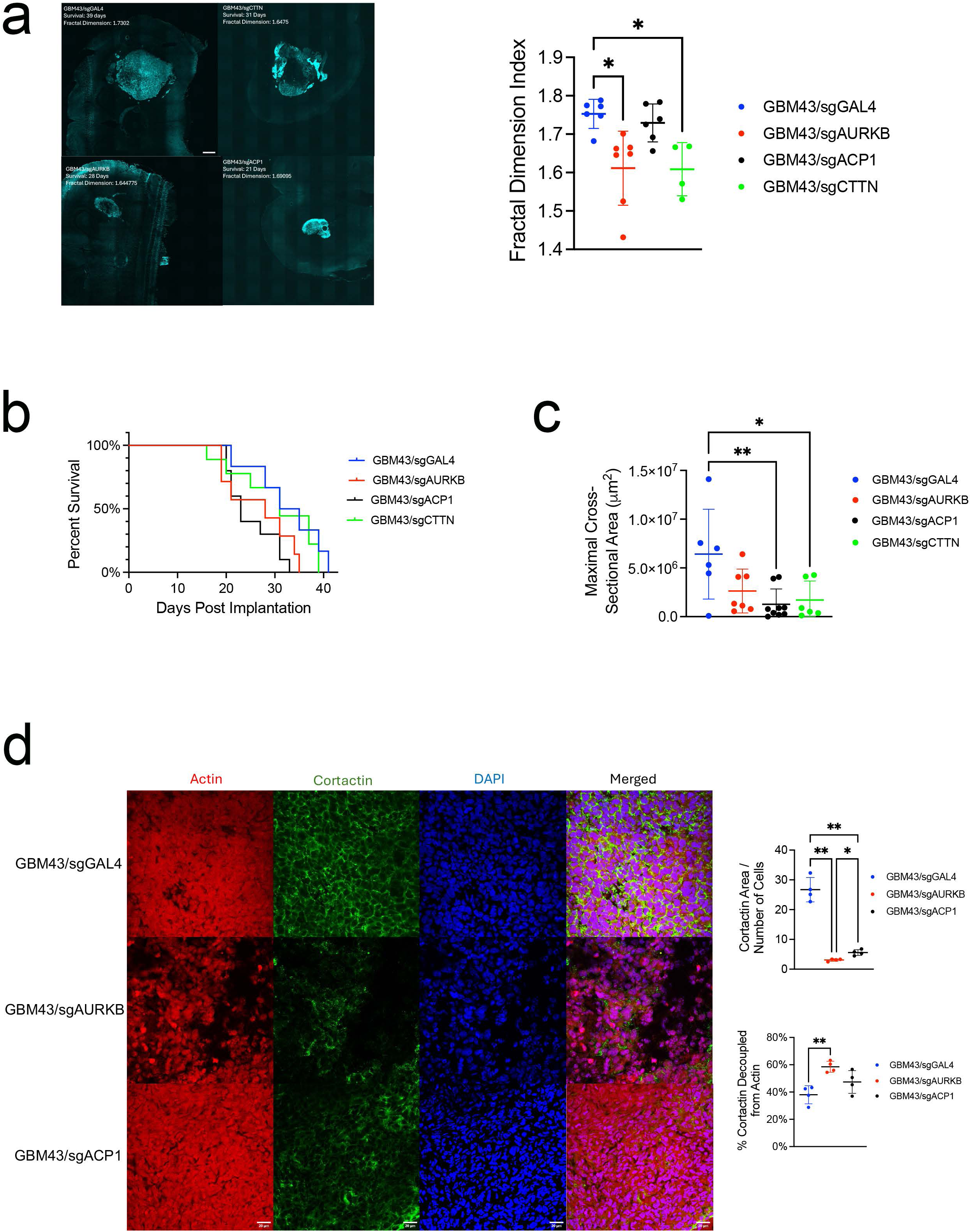
Targeting AURKB and ACP1 in orthotopic GBM xenografts *in vivo* slows invasion and uncouples cortactin from actin in tumor cells without affecting survival. GBM43/sgGAL4, GBM43/sgAURKB, GBM43/sgACP1, and GBM43/sgCTTN cells were implanted into the right frontal lobes of athymic mice and allowed to grow until mice reached endpoint, after which tumors were explanted and analyzed by immunofluorescence. **(a)** GBM43/sgAURKB (P=0.02; n=7) and GBM43/sgCTTN (P=0.02; n=4) xenografts were less invasive than GBM43/sgGAL4 (n=6) based on fractal analysis of images of tumors and their surrounding brain, which yields fractal dimension, a measure of invasive tumor growth pattern as a continuous number between 1 and 2, with higher numbers representing greater invasiveness. GBM43/sgACP1 (n=6) xenografts exhibited no change in fractal dimension compared to GBM43/sgGAL4 xenografts (P=0.8). Composite of images taken at 20x magnification, scale bar: 500 μm. **(b)** Knockdowns did not impact survival, with no difference noted in the survival of mice with GBM43/sgGAL4, GBM43/sgAURKB, GBM43/sgACP1, and GBM43/sgCTTN xenografts (P=0.6, n=6-10/group). **(c)** Maximal cross-sectional area of tumors at endpoint was noted to be smaller in GBM43/sgACP1 (n=9, P=0.007) and GBM43/sgCTTN (n=6, P=0.03) xenografts compared to GBM43/sgGAL4 (n=6) xenografts, but unchanged in GBM43/sgAURKB (n=7) xenografts compared to GBM43/sgGAL4 (P=0.08). **(d)** Immunofluorescence of xenografts at endpoint revealed decreased immunopositive cortactin area normalized to total area of cellular tumor in GBM43/sgAURKB (P=0.003) and GBM43/sgACP1 (P=0.005) xenografts relative to GBM43/sgGAL4 xenografts (upper graph) and increased decoupling of cortactin from actin in GBM43/sgAURKB (P=0.004) xenografts (lower graph). n=4/group, 60x magnification, scale bar: 20 μm. *P<0.05; **P<0.01; ***P<0.001; ****P<0.0001. Scatter dot plots show the mean (horizontal bar) with standard deviation (vertical bar).

We then used immunofluorescence to localize cortactin and actin in xenografts with *AURKB* or *ACP1* knockdown. Consistent with our results in cultured cells, *AURKB* and *ACP1* knockdown lowered intracellular cortactin levels in GBM43 xenografts (**Fig. 6d; Supp.** Fig. 5a), with *AURKB* or *ACP1* knockdown not altering intracellular cortactin localization (**Supp.** Fig. 5b). We also found that xenografts with *AURKB* knockdown, but not those with *ACP1* knockdown, exhibited disrupted intracellular actin-cortactin colocalization compared to xenografts with *GAL4* control targeting (**Fig. 6d**), aligning with the fractal index analysis results which were also impacted with *AURKB*, but not *ACP1*, knockdown.

### AURKB and ACP1 mediate distinct aspects of GBM invasion

While we had demonstrated shared functionality between AURKB and ACP1 in cultured GBM cells, such as stabilizing cortactin to promote its colocalization with actin, promotion of actin polymerization, and rendering of morphologic alterations conducive to GBM cell invasion, the distinct effects these proteins exerted *in vivo* with ACP1 impacting tumor size and AURKB impacting fractal index and actin-cortactin colocalization, prompted us to look for aspects of GBM cell invasion impacted differently by these two proteins, particularly in light of the potent anti-invasive effects seen when targeting both compared to either alone in culture (**Supp Fig. 1d**).

We began by looking at adhesion to ECM, the critical first step in the invasive process. We found that GBM43 cells with AURKB knockdown exhibited decreased adhesion to fibronectin, a critical component of the GBM ECM (*23*), while GBM43 cells with ACP1 knockdown had unchanged adhesion (**Fig. 7a**). We also found that GBM43 cells with AURKB, but not ACP1, knockdown exhibited increased individual cell stiffness as measured by atomic force microscopy (AFM) (P=0.0006; **Fig. 7b**).

**Figure 7.**
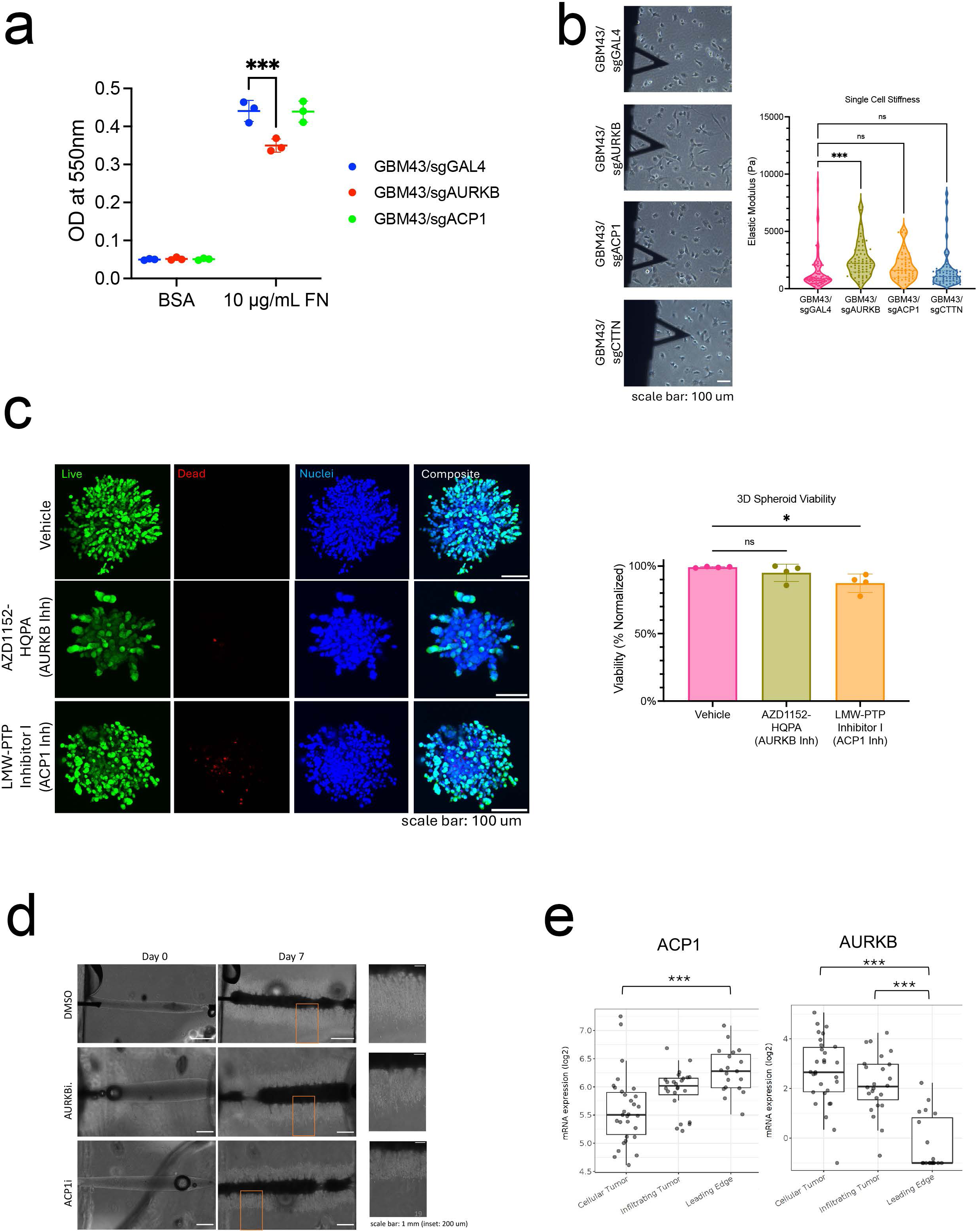
AURKB and ACP1 exert distinct effects on different aspects of GBM invasion. **(a)** GBM43/sgAURKB cells exhibited reduced adhesion to fibronectin compared to GBM43/sgGAL4 (n=3/group; P=0.0001). **(b)** AFM revealed decreased single-cell stiffness in GBM43/sgAURKB cells compared to GBM43/sgGAL4 cells (P=0.0006) with no changes noted in GBM43/sgACP1 or GBM43/sgCTTN cells (n=50-75 individual cells per condition). 10x, scale bar=100 µm. **(c)** Live/dead cell staining revealed that inhibition of ACP1 with LMW-PTP Inhibitor I led to decreased cell viability/increased cell death in GBM43 cells attempting to invade during neurosphere assays (P=0.04), with no changes noted when treating with AURKB inhibitor AZD1152-HQPA (n=4 spheres per condition). 10x, scale bar=100 µm. **(d)** Light microscopy revealed that, in U87 GBM cells, inhibition of ACP1 with LMW-PTP Inhibitor I caused alteration of the leader/follower dynamics we previously identified in 3D hydrogel assays, with no changes noted upon treatment with AURKB inhibitor AZD1152-HQPA. 10x, scale bar=1 mm (inset scale bar=200 µm). **(e)** *ACP1* expression increased from the core cellular tumor to the infiltrating tumor to the leading edge (P=4.4X10^-6^ cellular tumor vs. leading edge), while *AURKB* expression decreased from the core cellular tumor to the infiltrating tumor to the leading edge (P=4.4X10^-13^ cellular tumor vs. leading edge; P=4.9X10^-9^ infiltrating tumor vs. leading edge) as assessed using the Ivy Glioblastoma Atlas Project which transcriptomically profiled site-directed biopsies from these regions in newly diagnosed GBMs from 10 patients. *P<0.05; **P<0.01; ***P<0.001; ****P<0.0001

In contrast, GBM43 cells treated with LMW-PTP Inhibitor I to target ACP1, but not those treated with AURKB inhibitor AZD1152-HQPA, exhibited cell death in core cells attempting to invade in 3D neurosphere assays (P=0.04; **Fig. 7c**), suggesting a role for ACP1 in GBM cells migrating out of the tumor core. To corroborate the role for APC1 in GBM cell launch in a different context, we investigated its role in leader/follower dynamics we previously identified in 3D hydrogel assays (*24*). High molecular weight (HMW) gels were constructed (**Supp.** Figs. 6a-b) (*24*) to yield this previously described leader/follower phenotype. Interestingly, pharmacological inhibition of ACP1 with LMW-PTP Inhibitor I, but not AURKB with AZD1152-HQPA, in U87 GBM cells disrupted these leader/follower mechanics (**Supp.** Fig. 6c; **Fig. 7d**).

Together, these findings suggested that AURKB plays roles in the ability of GBM cells to anchor themselves and regulate their stiffness, functions attributed to myosin motors (*25*) and focal adhesions, while ACP1 plays roles in the ability of cells to withstand external stressors and launch out of the core. Analysis of human GBM gene expression data from Ivy GAP revealed that AURKB expression decreased during the transition from the core to the edge of patient GBMs, while ACP1 expression increased during this transition (**Fig. 7e**), supporting a role for the former in earlier stages of the invasive process and a role for the latter in later stages.

## DISCUSSION

While the hallmark of GBM and a defining contributor to its poor prognosis is invasion into the surrounding white matter, progress in understanding targetable mediators of this invasion has been limited. To close this knowledge gap, we developed a bioengineered 3D hydrogel invasion platform for high throughput screening of invasion mediators. The spatially dissectible nature of our hydrogel-based invasion devices allows us to analyze tumor cells in the invasive front versus non-invasive core of these devices, which in turn enables us to perform CRISPR screens in GBM cells invading these devices. Here, we used this approach, which we previously employed to define mediators of the metabolic needs of invading GBM cells (*4*), to identify druggable mediators of GBM invasion. We found that AURKB and ACP1, two hits from a druggable genome CRISPR screen, converge based on their serine phosphorylation and tyrosine dephosphorylation of cortactin, an actin-binding protein.

AURKB is a serine/threonine protein kinase found in the cell nucleus and cytoplasm whose most well-established function is its role as part of the chromosomal passenger complex (CPC), a series of proteins involved in regulating chromatin remodeling during cell division (*26*). Dysregulation of AURKB is often associated with cancer progression and poor prognosis. In GBM, specific targeting of AURKB can overcome temozolomide resistance (*11*), while use of a pan-AURK (A and B-targeting) agent creates a more favorable lipid metabolism profile and leads to reduced invasion of GBM stem cells (*27*).

ACP1 is a low molecular weight phosphotyrosine phosphatase implicated in various cellular processes, including signal transduction and actin cytoskeleton remodeling. Actin polymerization, crucial for cell motility and structure, is a key process in cancer cell invasion, enabling the formation of cellular protrusions like lamellipodia and invadopodia, which are necessary for tissue penetration and metastasis. In normal cells like osteoblasts, studies have demonstrated ACP1’s role in modulating cell adhesion and migration through its impact on focal adhesion dynamics (*28*), while in malignant cells like colorectal cancer cells, ACP1 has been implicated in cell migration (*12*), a critical step in cancer metastasis.

The actin-binding protein cortactin has been shown to have higher expression in glioma specimens than in non-tumor brain tissue with expression positively correlating with glioma grade and targeting of cortactin shown to reduce size and persistence time of lamellipodia and migration and invasion in 2D assays (*29*). Our work builds upon this observation by demonstrating the effect of targeting cortactin on GBM invasion in 3D assays and *in vivo* and gaining insights into regulators of cortactin phosphorylation. Human cortactin can be phosphorylated on serine residues S-113, S-405, and S-418 as well as on tyrosine residues Y-421, Y-470, and Y-486 (corresponding to Y-421, Y-466, and Y-482 in mouse cortactin) by PAK1, ERK1/2, and Src kinases. Here, we add to this finding by demonstrating the ability of AURKB to phosphorylate these serine residues of cortactin and the ability of ACP1 to dephosphorylate these tyrosine residues of cortactin.

Interestingly, our findings imply that both the serine phosphorylation and tyrosine dephosphorylation of cortactin promote GBM cell invasion and its associated cytoskeletal remodeling. These potentially differential effects of serine vs. tyrosine phosphorylation on cortactin biology are consistent with other proteins in which serine vs. tyrosine phosphorylation affects proteins differently. In particular, there is precedent for actin-binding proteins other than cortactin to be affected differently by tyrosine versus serine phosphorylation. For example, tyrosine phosphorylation of synaptopodin promotes its interaction with calcineurin, leading to changes in cell morphology, while serine/threonine phosphorylation of synaptopodin can have different regulatory effects (*30*). For cortactin, the different effects of serine vs. tyrosine phosphorylation are less well understood, limited to one cell biology study in which serine-phosphorylated cortactin promoted the assembly of dendritic actin networks, while tyrosine phosphorylation of cortactin appeared to regulate focal adhesion dynamics (*31*). Further work will be needed to determine the precise cellular effects of serine vs. tyrosine cortactin phosphorylation and whether differences in these effects can explain our findings, which imply that serine phosphorylation and tyrosine dephosphorylation of cortactin drives GBM invasion through effects on cortactin stability leading to changes in actin remodeling.

While our CRISPR screen had a higher false positive rate than typically arises from CRISPR screens done using the highly specific dCas9-KRAB-MeCP2 transcriptional repressor that we used (*32*), this likely reflects the fact we used different complementary methods for the screen versus validation. We used 3D hydrogel devices for the screen given the ability of these devices to house the larger number of cells needed for the screen, the fact that invasive versus non-invasive cells could be separated in these devices, and the ability of these devices to be used over a longer time period. For validation, we used 3D spheroid invasion assays, allowing for shorter-term studies requiring a smaller number of cells. The use of these complementary methods added greater stringency to our screen results and increased our confidence as we moved to mechanistic studies based on these results.

Indeed, beyond their potential value in identifying novel therapeutic strategies, druggable genome screens have the added benefit of identifying novel mechanisms of biological processes. By performing a druggable genome screen to look for inhibitors of GBM invasion in our biomimetic 3D hydrogels, we were able to identify AURKB and ACP1 as key druggable regulators of GBM invasion that activate a shared downstream target, cortactin, whose ability to promote actin assembly was implicated in its invasion-promoting effects.

While the invasion-promoting effects of AURKB were preserved *in vivo*, the fact that targeting AURKB or ACP1 did not affect survival in a murine model underscores the challenges of translating strategies that disrupt the key process of GBM invasion into survival benefits. Further work will be needed to determine whether the lack of these survival benefits reflects limitations in mouse models where invasion exerts far less of an adverse effect on survival than in humans (*33*) or whether anti-invasive therapies will need to be combined with other approaches that target proliferation directly in order to target proliferative pathways that might upregulate as part of a “go or grow” hypothesis (*34*) when invasion is targeted.

## METHODS

### Cell Culture

GBM43, GBM102, and GBM28 PDX cells were provided by the Mayo Clinic, while U87MG cells were obtained from ATCC. PDX lines were cultured in DMEM (Thermo Fisher) supplemented with 10% (vol/vol) fetal bovine serum (Corning, MT 35- 010-CV), 1% (vol/vol) penicillin-streptomycin (Thermo Fisher) and 1% (vol/vol) Glutamax (Thermo Fisher, 35-050-061). U87MG cells were cultured in DMEM (Thermo Fisher) supplemented with 10% (vol/vol) fetal bovine serum (Corning, MT 35-010-CV), 1% (vol/vol) penicillin-streptomycin (Thermo Fisher), 1% (vol/vol) Sodium Pyruvate, and 1 % (vol/vol) MEM Non-essential Amino Acids (Thermo Fisher). Cells were harvested using 0.25% Trypsin-EDTA (Thermo Fisher) and passaged under 30 times. Cells were screened bimonthly for mycoplasma and validated every six months by Short Tandem Repeat (STR) analysis at the University of California Berkely or San Francisco Cell Culture Facilities. For drug-treatment studies, cells were cultured in varying concentrations of AZD1152-HQPA or LMW-PTP Inhibitor I (MLS-0322825 cmpd 23).

### 3D Hydrogels

#### Me-HA Synthesis

Hyaluronic Acid (HA) hydrogels were synthesized as described (*24, 35*). Methacrylic anhydride (Sigma-Aldrich, 94%) was used to functionalize sodium hyaluronate (Lifecore Biomedical, Research Grade, LMW: 66–99 kDa, HMW: 1.5 MDa) with methacrylate groups (Me-HA). The extent of methacrylation per disaccharide was quantified by ^1^H nuclear magnetic resonance spectroscopy (NMR) and was ∼90-100% for materials used in this study. To add integrin-adhesive functionality, Me-HA was conjugated via Michael Addition with cysteine-containing RGD peptide Ac-GCGYGRGDSPG-NH2 (Anaspec) at 0.5 mmol/l.

#### HA Hydrogel Rheological Characterization

Hydrogel stiffness was characterized by shear rheology via a Physica MCR 301 rheometer (Anton Paar) with 8-mm parallel plate geometry for γ=0.5% and f=1 Hz. For low molecular weight (LMW) gels: Frequency was controlled to 50-1 Hz for the frequency sweep at a constant strain (γ=0.5%), and the modulus saturation curve with time was obtained under oscillation with constant strain (γ=0.5%) and frequency (f=1 Hz). Gel solution temperature was controlled (T=37°C) with a Peltier element (Anton Paar) and the sample remained humidified throughout the experiment. For high molecular weight (HMW) gels, gels were incubated externally in humidified chamber at 37°C for 2 hours. Hydrogels were then trimmed to fit the 8 mm probe; a frequency sweep was performed as above to determine stiffness, and a stress relaxing test (performed at 15% strain for 5 minutes, measuring stress every 0.5 s) was used to determine the stress relaxation percentage.

#### Tumorsphere Invasion Assays

Tumorspheres were fabricated using Aggrewell Microwell Plates (Stemcell Technologies). Briefly, 1.2x10^5^ cells were seeded into a single well of the Aggrewell plate to form spheroids with 100 cells. After 48 hours, spheroids were resuspended in phenol red-free serum-free Dulbecco’s Modified Eagle’s Medium (DMEM, Thermo Fisher, 21-063-029) at 1.5 spheroids/μL and used as solvent for HA hydrogel crosslinking. To form hydrogels, 6 wt.% Me-HA was crosslinked in phenol red-free serum-free Dulbecco’s Modified Eagle’s Medium (DMEM, Thermo Fisher, 21-063-029) with a protease-cleavable peptide (KKCG-GPQGIWGQ-GCKK, Genscript). HA-RGD gels were crosslinked with peptide crosslinkers at varying ratios to yield hydrogels with a shear modulus ∼300 Pa and a final 1.5 wt.% Me-HA (**Supp.** Fig. 1b). After 1 hour crosslinking in a humidified 37°C chamber, cell culture media was added to hydrogels and, unless noted, replenished every 2 days. To analyze spheroid invasion assay results, spheroids were imaged every two days using Eclipse TE2000 Nikon Microscope with a Plan Fluor Ph1 ×10 objective. Images were acquired using NIS-Elements Software. For each spheroid, invasion was calculated as (A_f_–A_i_)/A_i_) where A_f_=final spheroid area and A_i_=initial spheroid area. Spheroid area was measured in ImageJ and invasion was normalized to control spheroids.

#### Invasion Devices

To fabricate large invasion devices, the device base, lid, and spacers were laser-cut out of 1.5 mm thick CLAREX° acrylic glass (Astra Products). Pieces were assembled and fastened with epoxy, UV-treated for 10 min, and stored in cold room. On the day of experiment, devices were brought to room temperature and a 22GX1.5” bevel needle (BD Precision Glide) was inserted into the device as a channel mold. The HA hydrogel solution was casted around the wire and incubated for one hour in a humidified 37°C chamber. After crosslinking, devices with hydrogels and needles were submerged in culture media for at least 10 min, before removing the needle which left an open channel. Afterwards, four million cells were seeded into the open channel and channel ends were plugged with vacuum grease. Unless stated, devices were cultured for 28 days and media was replenished every 3 days, with 28 days chosen based on experiments revealing it to be when control GBM43 cells fully invade through hydrogels. Small devices were manufactured as previously described (*36*). Briefly, the device base and lid were laser cut out of 1.5 mm thick acrylic. The devices were assembled appropriately, and needle threading wires were threaded to provide the channel structure (Thomas Scientific). PDMS (Ellsworth Adhesive, 184 SIL ELAST KIT) was poured and allowed to cool: channels were then cut into the PDMS and the laser cut top was glued. Prior to seeding, small devices were treated with UV light for 20-30 min.

### CRISPRi Knockdown Screen

A lentiviral plasmid containing a dCAS9/KRAB/MeCP2 cassette was obtained from Addgene. Lenti-X 293T cells were transfected using this plasmid, after which virus containing the plasmid was generated and appropriate titers were determined. GBM43 cells were then transduced with the virus and selected using 5 mg/mL blasticidin to obtain a pure dCAS9/KRAB/MeCP2 positive population. To validate the integration of our dCAS9/KRAB/MeCP2 system, a sample of positive GBM43 cells were transduced with a plasmid containing an sgRNA against *ITGB1*. Cells were then harvested after 48h and knockdown efficiency was measured using a western blot.

Druggable genome sgRNA libraries were designed and screen performed as described (*37*). The druggable gRNA library consists of 13,250 sgRNAs against 2550 genes (5 sgRNAs/gene) and 250 sgRNAs against 50 genes acting as negative controls. Oligonucleotides for sgRNAs were synthesized by IDT, amplified by PCR, and cloned into pLG1 library vector which contains RFP as a selection marker. Lenti-X 293T cells were transfected using this cloned gRNA library plasmid. Virus containing the plasmid was then be generated and titrated appropriately. The sgRNA library was introduced into the GBM cells as a lentiviral pool. GBM43 cells expressing the CRISPRi machinery (dCAS9/KRAB/MeCP2) were infected with the virus and selected for RFP using flow cytometry. To ensure that only one sgRNA was introduced in a cell, viral infection was optimized to obtain 30-40% transfection efficiency.

To study differences in invasive capacity, we incorporated 25 million GBM cells expressing this library into 3D HA-RGD hydrogels. The top and bottom 10% of the most and least invasive cells, as measured by distance of invasion through the 3D matrix, were visualized using fluorescent imaging for RFP expression. Invasive cells were defined as those invading a distance greater than 200 µm from the channel wall. Genomic deoxyribonucleic acid (gDNA) was extracted from harvested cells using Monarch Genomic DNA Purification Kit (New England BioLabs, T3010S) per manufacturer’s protocol. gDNAs from six invasion devices were pooled and amplified by PCR, and sequenced using Illumina HiSeq 4000 for next-generation sequencing. Sequenced reads were aligned to the gRNAs in the CRISPR library and were quantified using the Screen Processing pipeline (https://github.com/mhorlbeck/ScreenProcessing). Appropriate negative controls were incorporated for the screen including non-functional and non-targeting gRNAs. Sequencing counts from samples were summed, normalized (count/million), and analyzed as single conditions. Fitness scores for each guide were calculated as log_2_ ratio of normalized counts. The median of the guides was used as the fitness score for each gene, and t-test assessed whether guides significantly deviated from 0.

### TurboID and Proteomics

The biotinylation of proximal proteins of ACP1 and AURKB was performed via lentiviral transduction using a TurboID vector C1(1–29)-TurboID-V5_pLX304 (Addgene 107175). The coding sequences for the amino acids of ACP1 and AURKB were cloned onto this vector using Gibson Assembly (NEB Gibson Assembly Master Mix, E2611S). The protein sequences of ACP1 and AURKB were PCR amplified from expression plasmids obtained from Origene (ACP1, CAT#: RC212653; AURKB, CAT#: RC210288). Cells transduced with TurboID were subjected to 10 µg/ml blasticidin selection. Prior to collection, GBM43 cells were treated with 10 µM biotin and incubated at 37°C for 10 minutes. Protein lysates were prepared using RIPA buffer, and after calibrating protein concentrations, streptavidin magnetic beads (Thermo Fisher, 88816) were added to the lysates and incubated at room temperature for one hour to allow bead-protein conjugation without introducing unspecific binding. Following the washing protocol (RIPA, KCL, Na_2_CO_3_, and Urea) (*38*), eluted proteins from the beads were collected for western blot analysis. Protein lysates were verified using antibodies against the V5 protein tag and biotin, and the streptavidin beads were sent for iTRAQ/TMT based mass spectrometry analysis. Peptide counts from mass spectrometry were analyzed for differential expression by DESeq2 package in R. Unique protein peptides are defined as enriched when compared to control if their log2Foldchange is larger than 1.

### Immunofluorescence Staining of Cultured Cells

GBM43 cells were cultured on coverslips coated with 1 µg/ml fibronectin in PBS for 4 hours and placed in 6-well plates. Once cells reached 50% confluency, they were fixed in 3.7% formaldehyde and 0.1% Triton X-100 in phosphate-buffered saline for 10 minutes. Fixed cells were stained with primary and secondary antibodies. Primary antibodies for ACP1 (R&D Systems, MAB5075-SP), AURKB (EnCor Biotechnology, MCA-6G2), cortactin (Proteintech, 11381-1-AP), phospho-Serine (Santa Cruz Biotechnology, sc-81514), phospho-Threonine (Santa Cruz Biotechnology, sc-5267) and p-Tyrosine (Santa Cruz Biotechnology, sc-7020) were used to characterize their distribution and quantity. DAPI was used to stain nuclei, and phalloidin (594 nm) was used for the actin cytoskeleton.

### Immunoblots and Immunoprecipitation

GBM43 cells, including GBM43/sgACP1 and GBM43/sgAURKB cells, were lysed in RIPA buffer supplemented with 10% protease inhibitor cocktail (Roche, # 05892970001) and 10% phosphatase inhibitor cocktail (Roche, # 4906845001). Lysates were quantified using the Pierce BCA kit (Thermo Fisher), and 10-30 µg of total proteins were used per loading. Protein samples were subjected to SDS-PAGE, denatured at 95°C, electrotransferred to membranes, and immunoblotted. For immunoprecipitation, 1000-3000 µg of cell lysate proteins were incubated with specific primary antibodies at 2 µg/ml overnight at 4°C. The immunoprecipitated complexes were verified by western blotting using a Licor Odyssey system.

### Gene Ontology Enrichment

For Gene Ontology (GO) enrichment analysis, we leveraged the Molecular Signatures Database (MSigDB) to collect KEGG gene set libraries and identify significantly enriched items. Initially, we compiled a list of genes whose encoded proteins were effective hits (log2FC > 1) for both ACP1 and AURKB. We then mapped these genes to their respective terms using annotations from the KEGG database. The KEGG library facilitated the enrichment analysis by calculating the enrichment scores for each gene set based on the frequency of genes within the term compared to a background set. Statistical significance was determined using Fisher’s exact test, and a Benjamini-Hochberg correction was applied to control for false discovery rates, with a threshold set at FDR < 0.01. Correlation of gene sets was computed using shared content genes before hierarchical clustering.

### Cytoscape

To visualize the networks of ACP1 and AURKB-affected targets that regulate key processes involved in cell invasion, we utilized Cytoscape, an open-source platform for complex network analysis and visualization. We imported the effective hits from TurboID proteomic data for ACP1 and AURKB, followed by performing gene ontology enrichment analysis on all genes. We then filtered the large global networks to focus on subsets of genes based on identified biological processes most relevant to cell invasion. CTTN, a hub gene among these processes, was highlighted, with nodes representing individual proteins and edges denoting their interactions. Visual attributes like node size were adjusted to reflect the enrichment of genes from TurboID proteomic data.

### Actin Polymerization Assay

Cell lysates were prepared in a non-denaturing form using IP lysis buffer (Thermo Fisher, 87787) and spun in 10kDa purification columns (Abcam, ab93349) at 10,000 rpm for at least 30 minutes. Lysate concentrations were measured with a BCA assay to ensure sufficient input (100 μg protein) for the actin polymerization assay. Reagents from the assay kit (Cytoskeleton, BK003) were reconstituted, and pyrene-labeled actin was snap-frozen in liquid nitrogen. All records were made in 96-well optical-bottom plates with solid polystyrene black upper structures and scanned using a Biotek synergy neo2 hybrid multimode reader. The fluorometer and kinetic measurement programs were set according to the product manual. Fluorescent signals over a 1-hour period were normalized to initial signals, and linear regression was performed to measure the slopes of the time courses.

### Tissue Processing and Sectioning

Tissue samples were harvested from mice that had reached experimental endpoint (15% loss of initial body weight) except in the case that humane endpoints, as designated by UCSF IACUC, were reached first. Cardiac perfusion was performed with PBS and whole brains were collected and immediately put on ice. Samples were transferred to 4% PFA for 24 hours, then placed into 15% (weight/vol) sucrose in PBS until infiltration occurred, then transferred to 30% sucrose (weight/vol). After equilibration, brains were embedded in OCT (Sakura Tissue Tech) and snap-frozen using a slurry of dry ice and 100% ethanol. Samples were stored at -80 for no more than one week before sectioning.

Samples were cross-sectioned on a cryostat at 10-micron width and transferred onto glass slides, with one sample present per slide, then stored at -80C until usage.

### Tumor Fractal Analysis

Unstained slides containing cross sections from the widest area of the tumor were allowed to thaw at room temperature for 10 minutes. Images were taken using the BC43 spinning disk confocal benchtop microscope at 20X magnification of the tumor’s entirety and the surrounding region of brain tissue, using the confocal blue channel to observe areas containing BFP, indicating tumor tissue. Images were then converted to binary and analyzed for fractal complexity based on Box Counting Dimension (D_B_) using the FracLac plugin on ImageJ.

### Tumor Tissue Immunostaining

Slides containing tumor sections were thawed at room temperature and fixed with 4% PFA for 10 minutes. Slides were blocked and permeabilized with 2.5% FBS (vol/vol), 2.5% BSA (weight/vol), 0.1% Triton-X-100 (vol/vol) in PBS for one hour. Antibody for cortactin staining (Proteintech Cortactin Rabbit PolyAb – Cat No. 11381-1- AP) was diluted 1:1000 to a final concentration of 650 ng/ml in blocking solution and applied to sections overnight at 4°C. Slides were then stained simultaneously with secondary antibody (Invitrogen Goat anti-Rabbit IgG (H+L) Cross-Adsorbed Secondary Antibody, Alexa Fluor^TM^ 488 and actin-staining reagent (Abcam Phalloidin iFluor 594), both diluted 1:1000 in blocking solution, for two hours in the dark at room temperature. Nuclear counterstaining was completed after secondary antibody staining, using NucRed^TM^ Dead 647 (TO-PRO-3 iodide) (Fisher) diluted to 2 drops/ml in PBS, for 20 minutes at room temperature. Slides were washed with PBS, air dried and mounted to coverslips with glycerol, and then stored in the dark at 4°C until imaging occurred. All images were taken via the BC43 spinning disk confocal microscope. The green, red, and far-red channels were used to image cortactin, actin, and nuclear counterstain respectively. Images were taken at 60x magnification throughout the tumor and ImageJ was used to subsequently complete quantitative IHC analysis.

### Adhesion Assay

Clear 96 well plates were coated with 100 μL of either 40 μg/mL Bovine Serum Albumin (Fraction V) (Fisher) or 10 ug/mL human fibronectin (Corning) in PBS and incubated overnight at 4°C. Cells were trypsinized, counted to 2.5 x 10^5^ cells/ml, and serum starved in DMEM without phenol red (Fisher) for 2 hours at 37°C.The plate was concurrently blocked using 0.5% BSA (weight/vol) in DMEM without phenol red for 1 hour at 37°C, then chilled on ice. 2 x 10^4^ cells in serum-free media solution were added to each well and the plate was incubated at 37°C for 90 minutes. Wells were then fixed with 100 μL of 4% paraformaldehyde (Fisher) for 15 minutes at room temperature, washed with PBS, then stained with Crystal Violet (Sigma Aldrich) for 10 minutes. The plate was washed and fully submerged in tap water, then left upside down to dry completely overnight. 50 μL of Sorenson’s Phosphate Buffer was added to each well the following morning and left to incubate at room temperature for 30 minutes. The optical density of each well was read at 550 nm via a BioTek Epoch Microplate Reader.

### Pharmacological Cell Viability

For 2D kill curves, cells were seeded at 20k cells/mL in a 48-well plate and allowed to adhere overnight. Wells were then treated with media and varying concentrations of the drug (in triplicate) and allowed to incubate for 48 hours. Cells were then fixed using 4 % PFA for 20 minutes and incubated with DAPI (1:1000) for 1 hour. Fluorescent images were taken on a Nikon Eclipse Ti-E epifluorescence microscope. Cells were counted using ImageJ, and total counts were normalized to the average number of cells in the vehicle condition to yield a percent viability value.

### 3D Live/Dead Assay

Cells were encapsulated following the spheroid protocol and allowed to invade for 3 days. Afterwards, cells were incubated with components from the Live/Dead Cell Imaging Kit (ThermoFisher, R37601) for 45 minutes. During the last ten minutes of incubation, Hoechst (ThermoFisher, H1399, 5 ug/mL) was added to visualize nuclei. Live, confocal fluorescence imaging was performed on a Zeiss LSM880 FCS confocal microscope. Using ImageJ, viability was quantified from the z-stack using the following formula: Viability (%) = 100*[(Area_total–_ Area_dead_)/ Area_total_], where Area_total_=Area_live_+Area_dead_.

### Hyperosmotic Pressure Assay

For the 2D assay, cells were seeded in 6-well plates and allowed to adhere overnight in normal media. Just prior to changing media, cell images were taken in 4x brightfield using a Nikon Eclipse Ti-E epifluorescence microscope Directly after imaging, media was removed and replaced with media containing 1.5 % v/v Poly-ethylene glycol (PEG) (TCI N0443). Cells were incubated for 3.5 hours and then imaged again. For the 3D assay, cells were seeded in 300 Pa hydrogels at 300 cells/μL and allowed to incubate overnight. Media was removed, and the PEG rich medium was added to half of the gels, and all gels were then incubated for another 24 hours. Gels were fixed with 4% PFA for 20 minutes and then incubated with DAPI (1:1000) and Wheat Germ Agglutinin (WGA, Invitrogen, W11262, 20 ug/mL) for 30 minutes. Fluorescent z stacks were taken using a Zeiss LSM880 FCS confocal microscope. Cells volumes were calculated in Fiji using the 3D Objects Counter plugin.

### Atomic Force Microscopy

Falcon Bacteriological Petri Dishes (Corning 351006) were briefly coated with Poly-D-Lysine hydrobromide (Millipore Sigma, 27964-99-4) dissolved in PBS (0.1 mg/mL) to assist with cell adhesion. Dishes were then washed with PBS for 3 minutes to remove residual poly-d-lysine. Cells were then plated at 100k cells per dish and allowed to adhere overnight. Cells were measured at 37°C under a HEPES-buffer medium with an MFP-3D-BIO AFM (Oxford Instruments) using pyramidal probes of nominal spring constant 0.08 N/m (PNP-TR-AU, Nanoworld) and a trigger threshold of 1 nN. Cantilevers were calibrated by the thermal method before using. Raw data was exported to MATLAB and Young’s modulus for each measurement was extracted by fitting the contact region to a pyramidal Hertz model with a Poisson ratio of 0.5.

### Statistics

Assays were done with at least three technical and biological replicates. To compare multiple groups, one-way ANOVA (parametric) or Kruskal-Wallis (non-parametric) tests were used for continuous outcome variables, with Chi-squared and Fisher Exact tests used for categorical outcome variables. ANOVA or Kruskal-Wallis tests were followed by Tukey or pairwise Wilcox post-hoc tests for comparisons between groups, respectively. Non-parametric two-tailed t-tests were used to compare two groups. Kaplan-Meier analysis was carried out for *in vivo* survival studies.

### Study Approval

Animal experiments were approved by UCSF IACUC (approval #AN105170-02).

## AUTHOR CONTRIBUTIONS

MH, AW, IL, EAA, and SJ designed the project, conducted experiments, processed data, interpreted results, and edited the manuscript. AL, AC, AR, AK, and ED conducted experiments, processed data, and interpreted results. SK and MKA procured funding, designed experiments, interpreted results, and wrote and edited the manuscript.

## ACKNOWLEDGEMENTS

A.W. was supported by the National Science Foundation Graduate Research Fellowship and the NIH T32 Biotechnology of Cell and Gene Therapy Predoctoral Fellowship. E.A.A was supported by the National Science Foundation Graduate Research Fellowship. S.K. was supported by R01CA227136, R01CA260443, and R01GM122375. M.K.A. was supported by R01CA227136, R01NS079697, and R01CA260443. We thank Mary West of the UC Berkeley Cell and Tissue Analysis Facility (CTAF). Work was performed in the QB3 CTAF that provided the Cryostar NX70. Confocal imaging occurred at the CRL Molecular Imaging Center, RRID:SCR_017852, supported by the Gordon and Betty Moore Foundation. We thank Holly Aaron and Feather Ives for microscopy advice.

