## Supplemental Figures 1-6 for "Druggable genome CRISPRi screen in 3D hydrogels reveals regulators of cortactin-driven actin remodeling in invading glioblastoma cells"

### SUPPLEMENTARY FIGURE LEGENDS

**Supplementary Figure 1. CRISPRi screen identifies druggable genes mediating GBM invasion in 3D hydrogels.**

**(a)** Schematic for how the druggable genome CRISPRi screen was done in GBM43 cells invading 3D HA-RGD hydrogels.

**(b)** Hydrogel storage modulus (G') (in Pascals) vs. time after the HA-RGD gel is casted. Time=0 represents the intersection of storage modulus (G') and loss modulus (G'') (n=3 hydrogels).

**(c)** Results of qPCRs confirming degree of knockdown relative to GBM43/sgGAL4. All knockdowns had P<0.001 for expression of the gene being targeted relative to GBM43/sgGAL4.

**(d)** Cell viability for GBM43 cells exposed to increasing concentrations of AURKB inhibitor AZD1152-HQPA or ACP-1 Inhibitor LMW-PTP Inhibitor I.

**(e)** Results of GBM43 spheroid invasion assay done in the presence of AURKB inhibitor AZD1152-HQPA, ACP1 inhibitor LMW-PTP Inhibitor I, or both inhibitors combined for 4 days (n=54-67 spheres per condition across 3 biological replicates). Scale bar: 100 $\mu$m.

**(f)** *Left*: Cell viability for GBM102 cells exposed to increasing concentrations of AURKB inhibitor AZD1152-HQPA and ACP1 inhibitor LMW-PTP Inhibitor I. *Right*: Results of neurosphere invasion assay for GBM102 cells grown in the presence of the highest nontoxic concentration of AURKB inhibitor AZD1152-HQPA (P<0.0001) or

ACP1 inhibitor LMW-PTP Inhibitor I (P=0.02) for 9 days (n=47-61 spheres across 3 separate biological replicates). Scale bar: 100 μm.

**(g)** *Left*: Cell viability for GBM28 cells exposed to increasing concentrations of AURKB inhibitor AZD1152-HQPA or ACP1 inhibitor LMW-PTP Inhibitor I. *Right*: Results of neurosphere invasion assay for GBM28 cells grown in vehicle vs. the presence of the highest nontoxic concentration of AURKB inhibitor AZD1152-HQPA (P=0.0004) or ACP1 inhibitor LMW-PTP Inhibitor I (P=0.02) for 11 days (n=32-49 spheres across 3 separate biological replicates). Scale bar: 100 μm.

*P<0.05; **P<0.01; ***P<0.001; ****P<0.0001.

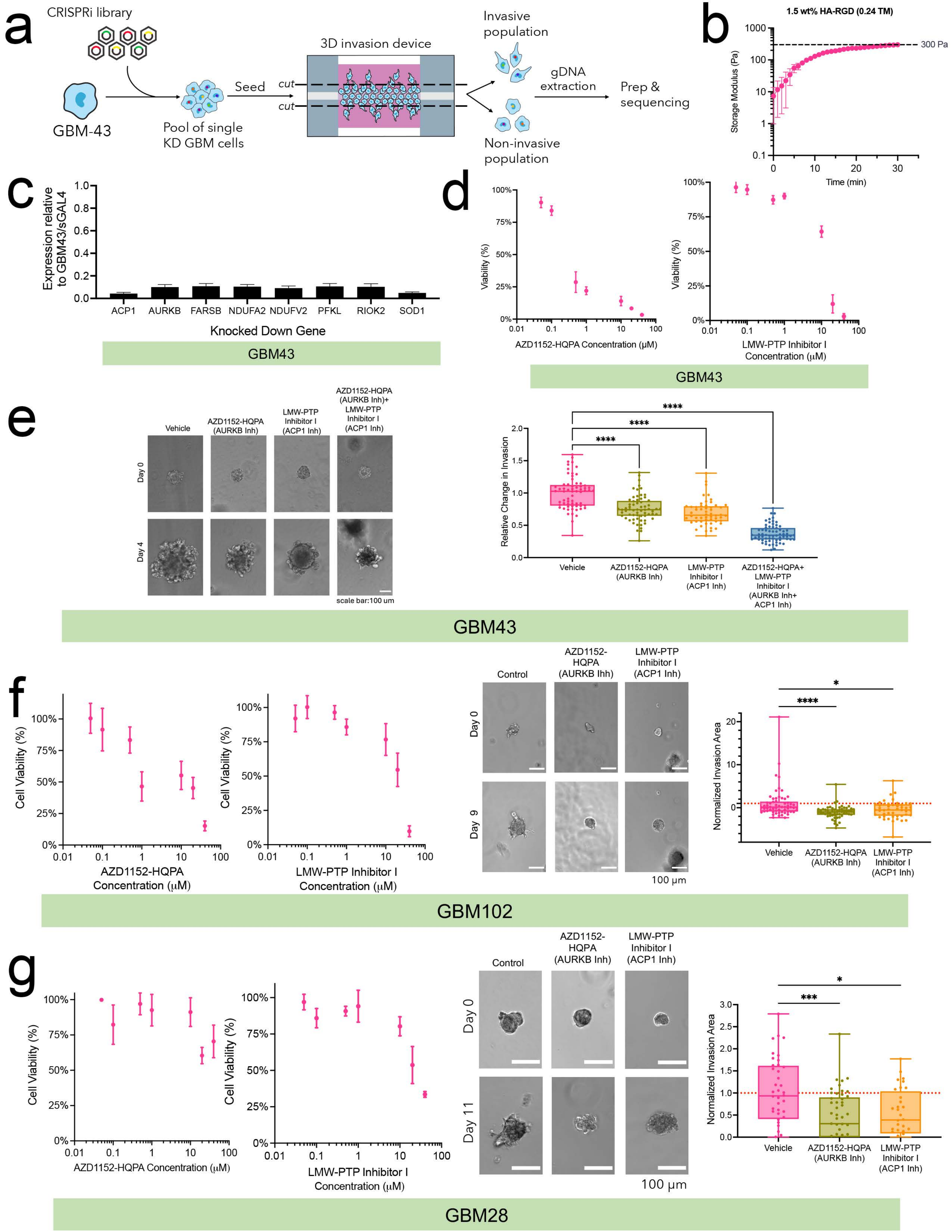
a
CRISPRi library
GBM-43
Pool of single KD GBM cells
Seed
3D invasion device
cut
cut
Invasive population
Non-invasive population
gDNA extraction
Prep & sequencing
b
1.5 wt% HA-RGD (0.24 TM)
Storage Modulus (Pa)
300 Pa
Time (min)
c
Expression relative to GBM43/sGAL4
ACP1
AURKB
FARSB
NDUFA2
NDUFV2
PFKL
RIOK2
SOD1
Knocked Down Gene
GBM43
d
Viability (%)
AZD1152-HQPA Concentration (µM)
LMW-PTP Inhibitor I Concentration (µM)
GBM43
e
Vehicle
AZD1152-HQPA (AURKB Inh)
LMW-PTP Inhibitor I (ACP1 Inh)
AZD1152-HQPA (AURKB Inh)+ LMW-PTP Inhibitor I (ACP1 Inh)
Day 0
Day 4
scale bar:100 um
Relative Change in Invasion
AZD1152-HQPA+ LMW-PTP Inhibitor I (AURKB Inh+ ACP1 Inh)
GBM43
f
Cell Viability (%)
AZD1152-HQPA Concentration (µM)
LMW-PTP Inhibitor I Concentration (µM)
Control
AZD1152-HQPA (AURKB Ihh)
LMW-PTP Inhibitor I (ACP1 Inh)
Day 0
Day 9
100 µm
Normalized Invasion Area
Vehicle
AZD1152-HQPA (AURKB Inh)
LMW-PTP Inhibitor I (ACP1 Inh)
GBM102
g
Cell Viability (%)
AZD1152-HQPA Concentration (µM)
LMW-PTP Inhibitor I Concentration (µM)
Control
AZD1152-HQPA (AURKB Inh)
LMW-PTP Inhibitor I (ACP1 Inh)
Day 0
Day 11
100 µm
Normalized Invasion Area
Vehicle
AZD1152-HQPA (AURKB Inh)
LMW-PTP Inhibitor I (ACP1 Inh)
GBM28

**Supplementary Figure 2. Proximity labeling identifies cortactin as a binding partner for AURKB and ACP1 with a role in GBM invasion and whose expression increases in invasive GBM cells.**

- **(a)** Results of qPCR confirming knockdown of *CTTN* in GBM43/sgCTTN relative to GBM43/sgGAL4 (n=3 replicates). P=0.007 for *CTTN* expression in GBM43/sgCTTN compared to GBM43/sgGAL4 cells.
- **(b)** Neurosphere invasion assay results in GBM43/sgCTTN relative to GBM43/sgGAL4. Scale bar: 100 μm; n=38-50 spheres/condition.

*P<0.05; **P<0.01; ***P<0.001; ****P<0.0001

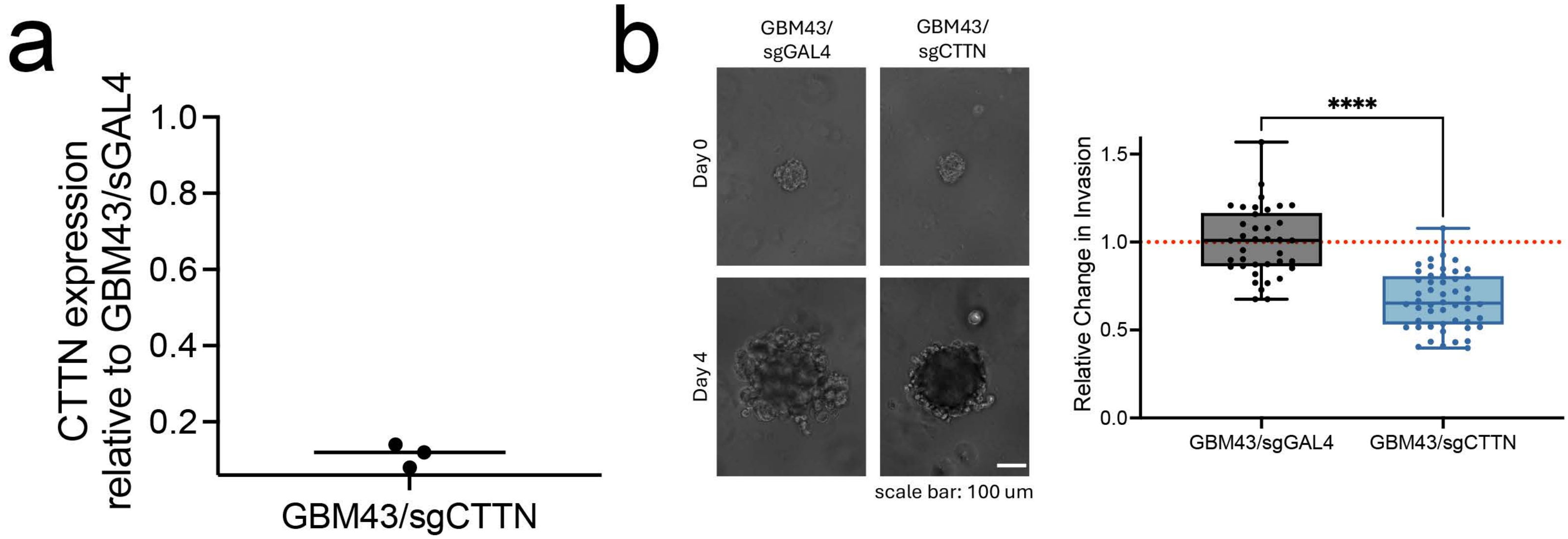
a
CTTN expression
relative to GBM43/sGAL4
1.0
0.8
0.6
0.4
0.2
GBM43/sgCTTN
b
GBM43/
sgGAL4
GBM43/
sgCTTN
Day 0
Day 4
scale bar: 100 um
****
Relative Change in Invasion
1.5
1.0
0.5
0.0
GBM43/sgGAL4
GBM43/sgCTTN

**Supplementary Figure 3. Serine phosphorylation of cortactin by AURKB and tyrosine dephosphorylation of cortactin by ACP1.**

**(a)** Shown are the three technical replicates of immunoprecipitation in which cortactin was immunoprecipitated from GBM43 cell lysates followed by blotting for phosphorylated serine/threonine (row one), blotting for cortactin (row two), and blotting the non-precipitated lysates for β-actin (row three).

**(b)** Shown are the three technical replicates of immunoprecipitation in which phosphorylated tyrosine was immunoprecipitated from GBM43 cell lysates followed by blotting for cortactin (row one) and blotting the non-precipitated lysates for cortactin (row two) and β-actin (row three).

a

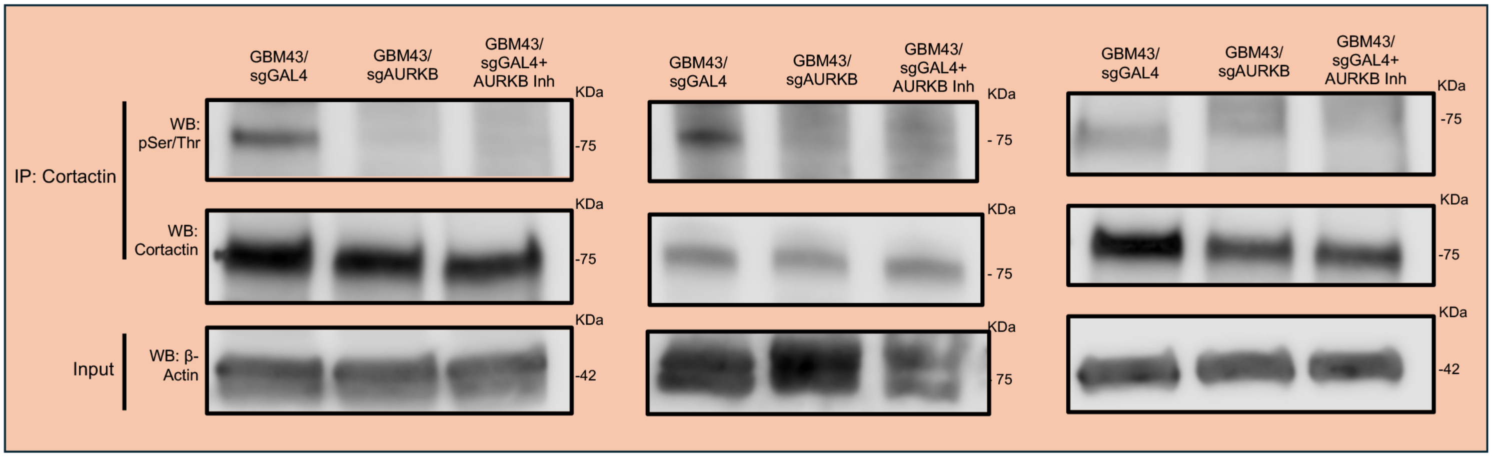
GBM43/ sgGAL4
GBM43/ sgAURKB
GBM43/ sgGAL4+ AURKB Inh
KDa
IP: Cortactin
WB: pSer/Thr
-75
WB: Cortactin
-75
Input
WB: β-Actin
-42
75

b

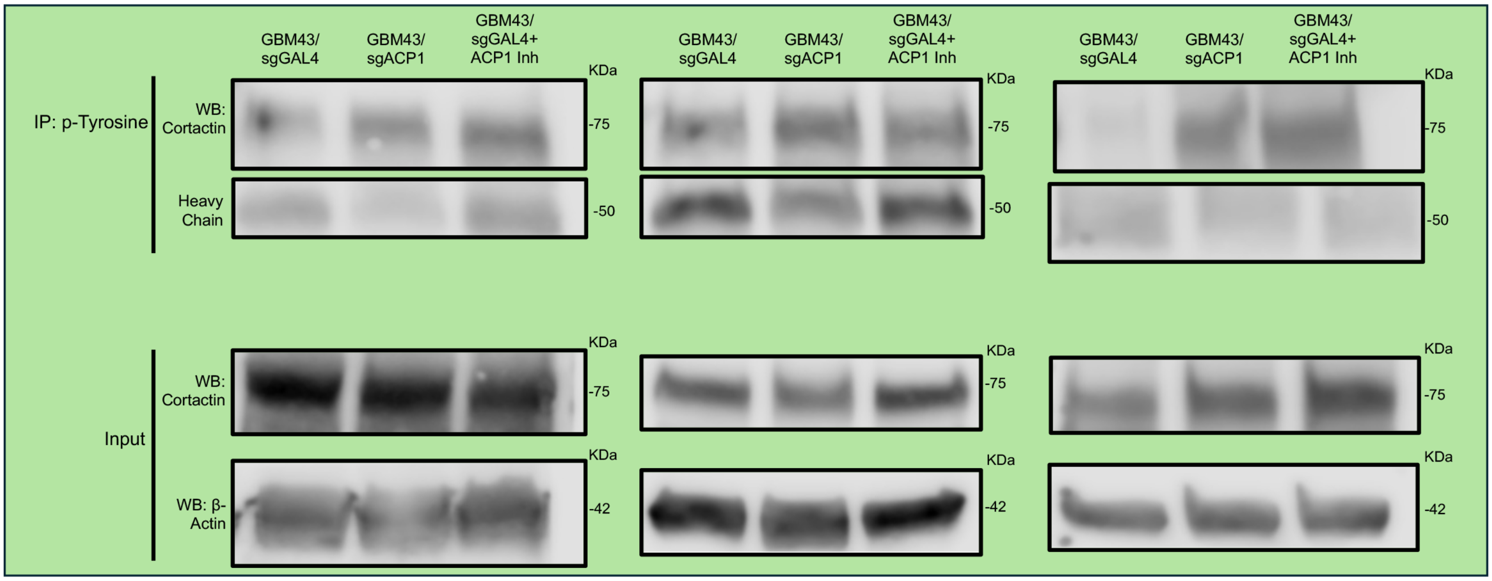
GBM43/ sgGAL4
GBM43/ sgACP1
GBM43/ sgGAL4+ ACP1 Inh
KDa
IP: p-Tyrosine
WB: Cortactin
-75
Heavy Chain
-50
Input
WB: Cortactin
-75
WB: β-Actin
-42

**Supplementary Figure 4. AURKB and ACP1 effects on cortactin phosphorylation alter actin polymerization and cell morphology.**

**(a)** Cell lysates from GBM43/sgCTTN (red) and GBM43/sgGAL4 cells (black) were used in actin polymerization assays, with GBM43/sgCTTN lysates slowing actin polymerization relative to GBM43/sgGAL4 lysates (assessed by comparing the slope of the polymerization curves derived from linear regression, $P<0.0001$). Differences in polymerization levels at individual timepoints did not occur during the assay but would be projected to eventually occur based on the different polymerization rates identified by slope comparison. n=3/group.

**(b)** Targeting AURKB and ACP1 in GBM43/sgAURKB and GBM43/sgACP1, respectively, creates a more circular cell morphology ($P<0.0001$) that is less conducive to invasion, as assessed by measuring the aspect ratio using images (not shown) taken of cells grown in two-dimensional culture. n=63-67 cells/group.

**(c)** *Left*: GBM43/sgAURKB ($P<0.0001$) and GBM43/sgACP1 ($P=0.002$) cells had smaller 3D volumes when placed into 3D HA-RGD devices than GBM43/sgGAL4 cells, while GBM43/sgCTTN cells were unchanged in size ($P=0.08$) (n=26-66 cells per condition). *Right*: GBM43/sgAURKB ($P=0.0001$), GBM43/sgACP1 ($P<0.0001$), and GBM43/sgCTTN ($P<0.0001$) cells had smaller 2D areas compared to GBM43/sgGAL4 cells (n=78-84 cells per condition) when placed into 3D HA-RGD devices.

**(d)** When exposed to hyperosmotic stress in the form of PEG, the contractile response occurring in GBM43/sgGAL4 cells ($P=0.0002$) did not occur in GBM43/sgAURKB, GBM43/sgACP1, or GBM43/sgCTTN cells likely due to their small starting size.

**(e)** Targeting AURKB with CRISPRi or drug (AZD1152-HQPA) in cultured GBM43 cells leads to reduced cortactin serine phosphorylation (based on overlapping area of cortactin and p-Ser/p-Thr staining) normalized to cell area in the total cell ($P<0.0001$ with CRISPRi or with drug), in the cell core ($P<0.0001$ with CRISPRi or with drug), and within 3 μm of the cell edge ($P=0.008$ with CRISPRi, $P=0.007$ with drug).

**(f)** Targeting ACP1 with CRISPRi or drug (LMW-PTP Inhibitor I) in cultured GBM43 cells leads to increased cortactin tyrosine phosphorylation (based on overlapping area of cortactin and p-Tyr staining) normalized to cell area in the total cell ($P=0.01$ with CRISPRi, $P=0.0005$ with inhibitor) and cell core ($P=0.01$ with CRISPRi, $P<0.0001$ with inhibitor), but with no change within 3 μm of the cell edge ($P=0.1$ with CRISPRi, $P=0.06$ with inhibitor) due to higher background cortactin tyrosine phosphorylation at the cell edge.

*$P<0.05$; **$P<0.01$; ***$P<0.001$; ****$P<0.0001$

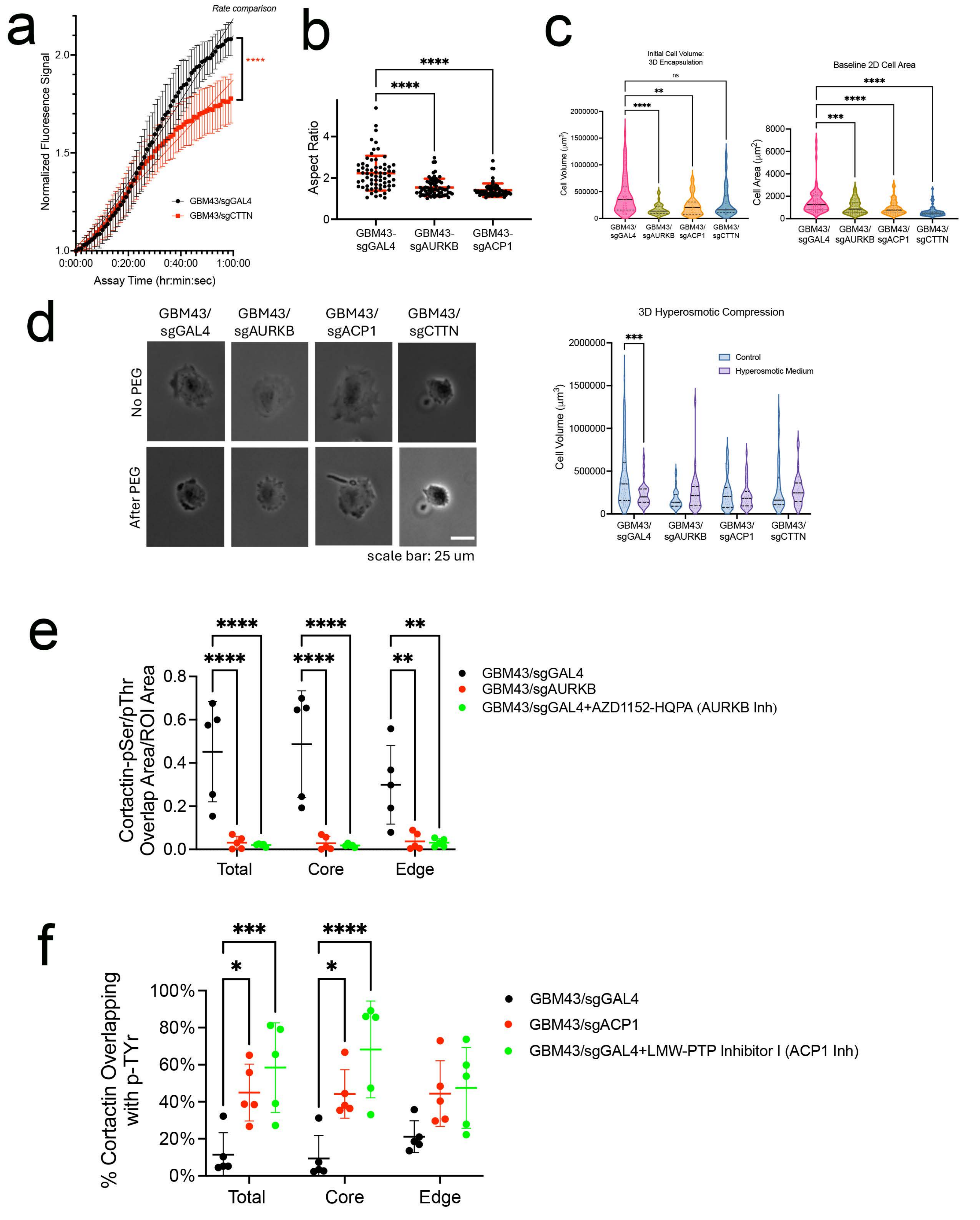

a
Rate comparison
Normalized Fluoresence Signal
Assay Time (hr:min:sec)
GBM43/sgGAL4
GBM43/sgCTTN
****
b
Aspect Ratio
GBM43-sgGAL4
GBM43-sgAURKB
GBM43-sgACP1
****
c
Initial Cell Volume: 3D Encapsulation
Baseline 2D Cell Area
Cell Volume (μm³)
Cell Area (μm²)
GBM43/sgGAL4
GBM43/sgAURKB
GBM43/sgACP1
GBM43/sgCTTN
ns
d
GBM43/sgGAL4
GBM43/sgAURKB
GBM43/sgACP1
GBM43/sgCTTN
No PEG
After PEG
scale bar: 25 um
3D Hyperosmotic Compression
Control
Hyperosmotic Medium
Cell Volume (μm³)
e
Cortactin-pSer/pThr Overlap Area/ROI Area
Total
Core
Edge
GBM43/sgGAL4
GBM43/sgAURKB
GBM43/sgGAL4+AZD1152-HQPA (AURKB Inh)
f
% Cortactin Overlapping with p-TYr
Total
Core
Edge
GBM43/sgGAL4
GBM43/sgACP1
GBM43/sgGAL4+LMW-PTP Inhibitor I (ACP1 Inh)

**Supplementary Figure 5. Targeting AURKB and ACP1 in orthotopic GBM xenografts *in vivo* uncouples cortactin from actin in tumor cells.**

**(a)** Immunofluorescence of xenografts at endpoint revealed decreased immunopositive cortactin area normalized to total area of cellular tumor in GBM43/sgAURKB (P=0.003, n=7) and GBM43/sgACP1 (P=0.005, n=6) xenografts relative to GBM43/sgGAL4 (n=6) xenografts, and virtually undetectable decreased immunopositive cortactin area normalized to cell number in GBM43/sgCTTN xenografts (n=4) relative to the other three groups (P=0.003 vs. GBM43/sgACP1, P=0.0007 vs. GBM43/sgAURKB, P=0.004 vs. GBM43/sgGAL4). Shown are images from **Fig. 6d** alongside images form GBM43/sgCTTN xenografts that were not included in **Fig. 6d**. 60x magnification, scale bar: 20 μm.

**(b)** Tumor cells in GBM43/sgGAL4, GBM43/sgAURKB, and GBM43/sgACP1 xenografts had comparable intracellular cortactin distribution (P=0.7-0.9), with the portion of cortactin at the cell edge unchanged across all 3 groups.

*P<0.05; **P<0.01; ***P<0.001; ****P<0.0001

a

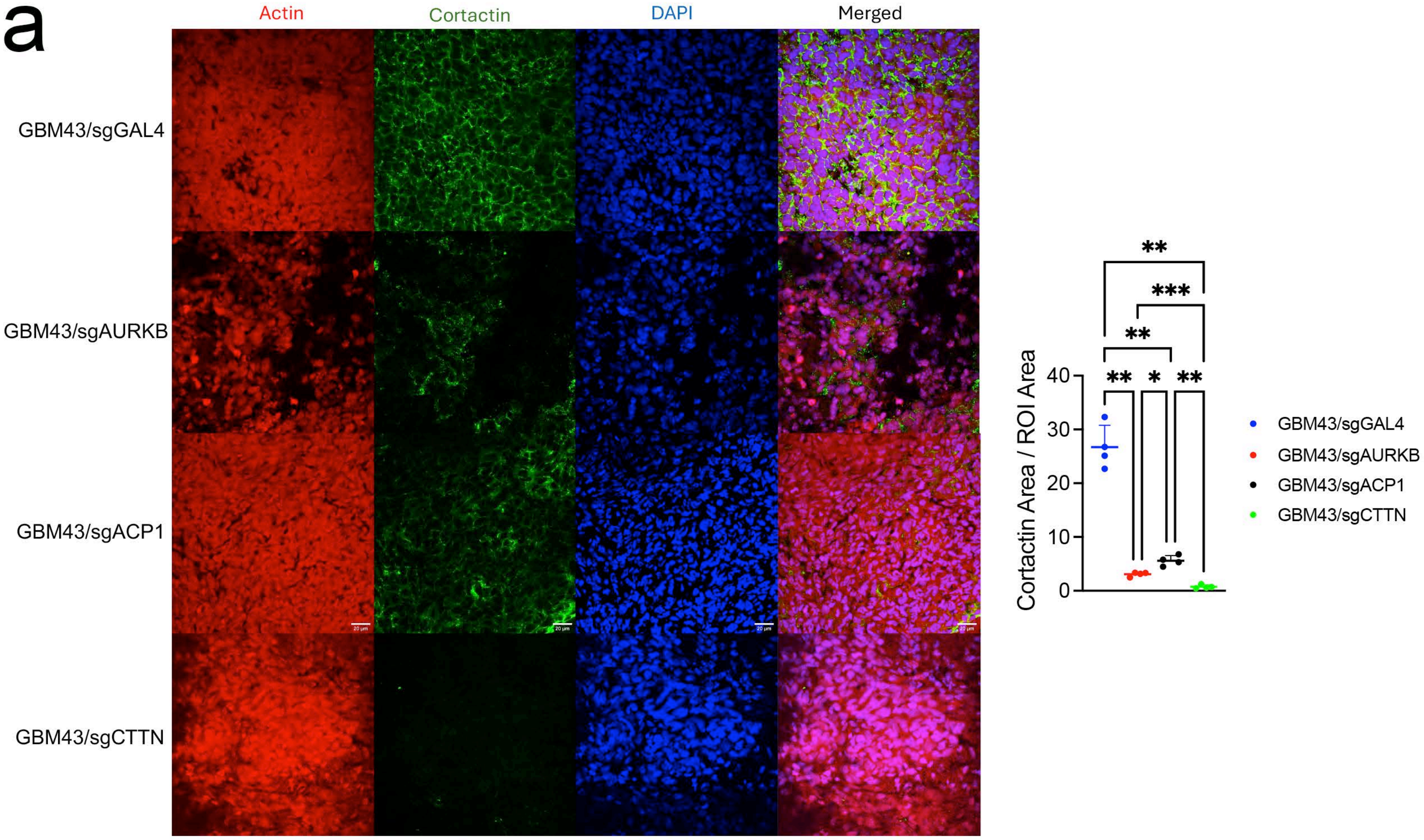
Actin
Cortactin
DAPI
Merged
GBM43/sgGAL4
GBM43/sgAURKB
GBM43/sgACP1
GBM43/sgCTTN
Cortactin Area / ROI Area
40
30
20
10
0
**
***
**
**
*
**
GBM43/sgGAL4
GBM43/sgAURKB
GBM43/sgACP1
GBM43/sgCTTN

b

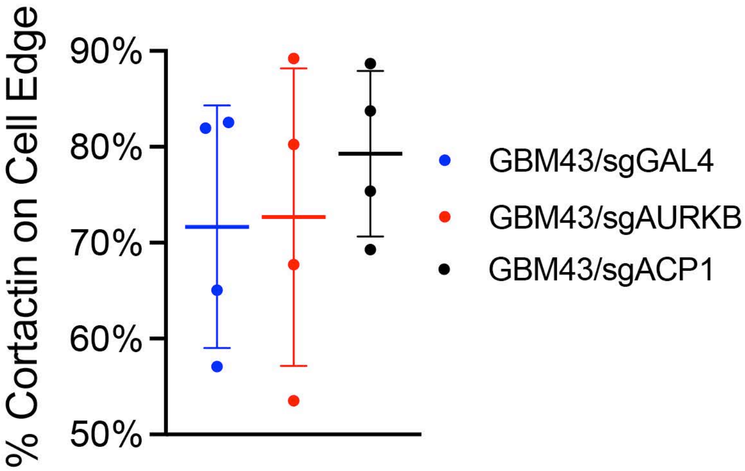
% Cortactin on Cell Edge
90%
80%
70%
60%
50%
GBM43/sgGAL4
GBM43/sgAURKB
GBM43/sgACP1

**Supplementary Figure 6. AURKB and ACP1 exert distinct effects on different aspects of GBM invasion.**

**(a)** Hydrogel storage modulus (G') (in Pascals) of HMW precast gels.

**(b)** Stress relaxation curve showing degree of stress relaxation of material. The normalized relaxation modulus is obtained by normalizing points at all times to the modulus at 0 seconds.

**(c)** U87 cell survival after treating cultured cells with varying concentrations of AURKB inhibitor AZD1152-HQPA (left) or ACP1 inhibitor LMW-PTP Inhibitor I (right) (n=3 biological replicates per concentration).

*P<0.05; **P<0.01; ***P<0.001; ****P<0.0001

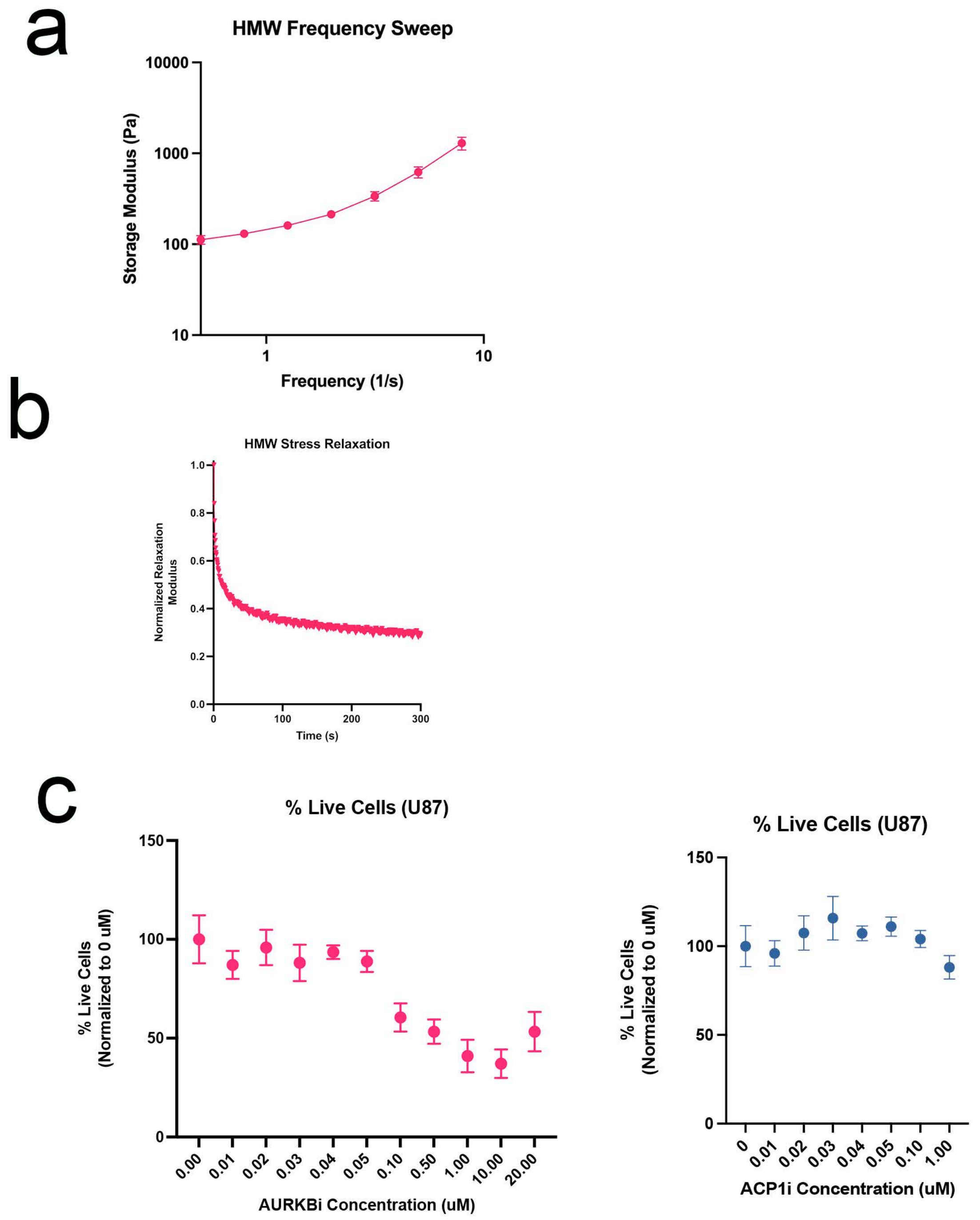
a
HMW Frequency Sweep
Storage Modulus (Pa)
10000
1000
100
10
1
10
Frequency (1/s)
b
HMW Stress Relaxation
Normalized Relaxation Modulus
1.0
0.8
0.6
0.4
0.2
0.0
0
100
200
300
Time (s)
c
% Live Cells (U87)
% Live Cells (Normalized to 0 uM)
150
100
50
0
0.00
0.01
0.02
0.03
0.04
0.05
0.10
0.50
1.00
10.00
20.00
AURKBi Concentration (uM)
% Live Cells (U87)
% Live Cells (Normalized to 0 uM)
150
100
50
0
0
0.01
0.02
0.03
0.04
0.05
0.10
1.00
ACP1i Concentration (uM)
